# CENP-N promotes the compaction of centromeric chromatin

**DOI:** 10.1101/2021.06.14.448351

**Authors:** Keda Zhou, Magdalena Gebala, Dustin Woods, Kousik Sundararajan, Garrett Edwards, Dan Krzizike, Jeff Wereszczynski, Aaron F. Straight, Karolin Luger

## Abstract

The histone variant CENP-A is the epigenetic determinant for the centromere, where it is interspersed with canonical H3 to form a specialized chromatin structure that nucleates the kinetochore. The arrangement of nucleosomes at the centromere into higher order structure is unknown. Here we demonstrate that the CENP-A interacting protein CENP-N promotes the stacking of CENP-A containing mono-nucleosomes and nucleosomal arrays through a previously undefined interaction between the α6 helix of CENP-N with the DNA of a neighboring nucleosome. We describe the cryoEM structures and biophysical characterization of such CENP-N mediated nucleosome stacks and nucleosomal arrays and demonstrate that this interaction is responsible for the formation of densely packed chromatin at the centromere in the cell. Our results provide first evidence that CENP-A, together with CENP-N, promotes specific chromatin higher order structure at the centromere.

**One-Sentence Summary:** The centromere-associated protein CENP-N promotes centromere-specific nucleosome stacking and higher order structures in vitro and in the cell.

---

The three-dimensional arrangement of nucleosomes (each consisting of two copies of histones H3, H4, H2A, and H2B that wrap 147 base pairs (bp) of DNA (*1*)) determines local and global chromatin architecture in all eukaryotes. While many *in vitro* studies provide evidence for a defined 30 nm fiber where nucleosomes are regularly packed through the interactions between n and n+2 nucleosomes, this has not been observed in the cell (*2, 3*). Instead, chromatin fibers are folded irregularly and diversely, with much variability between cell states and genome loci. Molecular dynamics simulations suggest that the energy barriers between different nucleosome arrangements are relatively low (*4*). Variations of nucleosome composition, such as DNA sequence, length of linker DNA connecting individual nucleosomes, incorporation of histone variants, and post-translational modifications of histones all have the potential to affect chromatin condensation and thus DNA accessibility, either directly or through the recruitment of a plethora of interacting factors (*5*).

The centromere is a specialized chromatin region onto which the kinetochore assembles. This megadalton >100 protein complex ultimately promotes faithful chromosome segregation by forming attachment points to the mitotic spindle. Nucleosomes containing the centromeric histone H3 variant CENP-A distribute among canonical nucleosomes in a ∼1:25 ratio (possibly in a clustered arrangement (*6*)), providing the sole epigenetic determinant of the centromere (*7*). The crystal structure of CENP-A-containing nucleosomes shows that rather than wrapping the canonical 147 bp of DNA, it stably binds only 121 bp DNA. This results in the unique organization of CENP-A containing tri-nucleosomal arrays (*8*), and renders the CENP-A nucleosome unable to bind linker histone H1 (*9, 10*). The key function of CENP-A nucleosomes appears to be the recruitment of centromere-specific proteins, most notably CENP-N and CENP-C, both of which recognize unique features of nucleosomal CENP-A, and upon which the CCAN complex and ultimately the kinetochore assemble (reviewed in (*11*)). Both CENP-N and CENP-C affect CENP-A nucleosome dynamics and structure *in vitro* at the mono-nucleosome level (*12–17*), but their effect on chromatin higher order structure has not been investigated at the molecular level (*18*).

## CENP-N promotes the stacking of CENP-A mono-nucleosomes

Previously, we reported the cryoEM structure of a CENP-A nucleosome reconstituted with the 601 nucleosome positioning DNA in complex with the nucleosome-binding domain of CENP-N (CENP-N^1-289^) (*13*). In this structure, CENP-N makes extensive contacts with DNA through the pyrin domain and the CNL-HD (CENP-N-CENP-L homology domain) and specifically recognizes the CENP-A histone through the α1 helix and β3-β4 loop of CENP-N (*13*). α-satellite DNA is more typical of the DNA sequence at the centromere, and we therefore determined the cryoEM structure of the CENP-A nucleosome, reconstituted onto palindromic α-satellite DNA (*1*), bound to CENP-N^1-289^, to a resolution of 2.7 Å (fig. S1). These structures are very similar to previously published structures on 601 DNA (*13, 19, 20*); see additional analysis and discussion of the higher-resolution structure in supplementary information (fig. S14).

In the cryoEM images we consistently observed that ∼30% of CENP-A nucleosomes formed ordered stacks on the grid in presence of CENP-N, ranging from 2-10 nucleosomes (fig. S1A, S2). CENP-A nucleosomes reconstituted with the 601 nucleosome positioning sequence (*21*) exhibit the same behavior in the presence of CENP-N (fig. S3A). By focusing single particle analysis on nucleosome pairs contained within these stacks, defined density for CENP-N was observed between two nucleosomes in the 2D class averages for nucleosomes reconstituted on either DNA fragment (fig. S2B and S3B). After 3D reconstruction and refinement (fig. S2A, S3C), we obtained cryoEM maps in which two or one CENP-N^1-289^ could be unambiguously docked between two CENP-A nucleosomes (fig. 1A, shown for α-sat nucleosomes). Superposition of 3D maps (fig. S3D) showed the exact same nucleosome stacks in cryoEM datasets of CENP-N with CENP-A nucleosomes reconstituted onto the two different DNA sequences, suggesting that DNA sequence context does not affect nucleosome stacking. As such, all following experiments were performed with 601 nucleosomes.

**Fig. 1.**
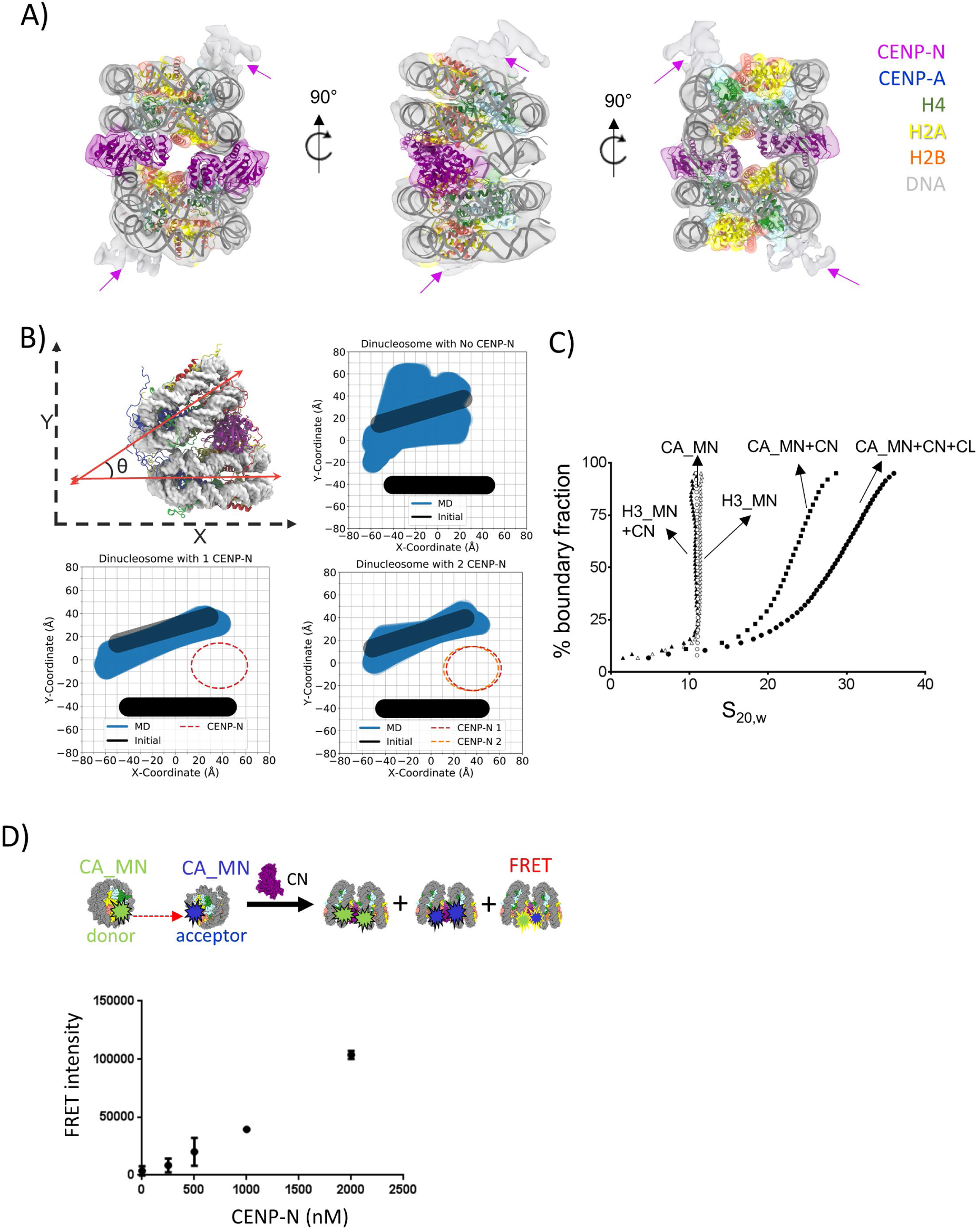
CENP-N mediates CENP-A nucleosome stacking *in vitro*. **A)** A model of two CENP-A nucleosomes (α satellite DNA) connected by two copies of CENP-N was fit into the density map. Protein identity is indicated by color codes. Arrows highlight the weak density attributed to a second CENP-N on the other side of the CENP-A nucleosomes. **B)** Simulation plots of nucleosome stacks in the absence or presence of CENP-N. This plot used two coordinate points from each nucleosome, namely the geometric center of the C1’-atoms from nucleotides located at the dyad and opposite of the dyad (fig, S4 for illustration). The C1’ atoms of every nucleotide in one nucleosome, pictured as the bottom nucleosome of each graph in figure 1B, were used to align each frame of the trajectory. The dyad and opposing points in the other nucleosome, pictured as the top nucleosome of each graph in figure 1B, were plotted to depict the relative sampling of each stacked-nucleosome system. **C)** SV-AUC (Enhanced van Holde-Weischet plots) for CENP-A (CA) or H3 mono-nucleosomes (MN) in complex with CENP-N^1-289^ (CN) or full length CENP-N/CENP-L (NL), as indicated in the plot. **D)** FRET analysis of CENP-A mono-nucleosome (CA_MN) interactions in the absence or presence of CENP-N. Donor is CENP-A mono-nucleosomes containing Alexa 488 labeled H2B; Acceptor is a CENP-A mono-nucleosome containing Atto N 647 labeled H2B. 250 nM donor and acceptor nucleosome; FRET intensity in dependence of [CENP-N]. Error bars from four independent measurements of two biological replicates.

Molecular dynamics (MD) simulations were performed to evaluate the stability of CENP-N binding to nucleosomes in solution and how it influences the dynamics of di-nucleosomal stacking (movie S1-S3). By plotting a cross-sectional view of the nucleosome coordinates as seen in figure 1B, we were able to determine the relative stabilizing effect provided by each CENP-N to stacked nucleosomes. The stacking of two mono-nucleosomes is rather unstable in simulations without CENP-N, where the nucleosomes explores a wide range of relative orientations. Stacked nucleosomes exhibit a similar amount of sampling whether they are in complex with one or two copies of CENP-N (fig. 1B). Other metrics for the inter-nucleosomal interactions, including the relative rise, shift, and tilt, exhibited similar trends (fig. S4), suggesting that a singular CENP-N is sufficient to stably maintain stacking between two CENP-A nucleosomes.

To confirm that nucleosome stacks are not artifacts of cryoEM grid preparation, we analyzed CENP-A nucleosomes in the absence and presence of CENP-N by sedimentation-velocity analytical ultracentrifugation (SV-AUC) under the buffer conditions used for CryoEM, but at much lower nucleosome concentrations (250 nM compared to the µM concentrations required for cryoEM). In the absence of CENP-N, CENP-A mono-nucleosomes sediment homogenously with a sedimentation coefficient (S_(20,W)_) of ∼10.5 S (fig. 1C), consistent with reported values for canonical nucleosomes (*22*). In the presence of CENP-N^1-289^, CENP-A nucleosomes assemble into much larger and more heterogeneous species, as evident by S_(20,W)_ ranging from 13 S to 30 S. For reference, dinucleosomes and 12mer-nucleosomal arrays sediment at 13 S and 30 S, respectively (unpublished results and (*23*)). When CENP-N was combined with nucleosomes containing H3, no larger species were observed upon addition of CENP-N (fig. 1C). To analyze the effect of CENP-N in a more physiologically relevant context, we showed that full length CENP-N in complex with CENP-L bound to a CENP-A nucleosome under the same conditions also promotes the oligomerization of CENP-A nucleosome (fig. 1C). To further confirm that nucleosomes indeed come in close contact in the presence of CENP-N, we designed a Foerster resonance energy transfer (FRET) assay. CENP-A nucleosome containing Alexa 488 labeled H2B was the donor, and CENP-A nucleosome containing Atto N 647 labeled H2B served as the acceptor. Nucleosome-nucleosome interactions should result in a strong FRET signal, and indeed we observed an increase in FRET upon titrating CENP-N into an equimolar mixture of donor and acceptor nucleosomes (fig. 1D). Collectively, our data shows that CENP-N mediates the stacking of mono-nucleosomes by engaging simultaneously with two CENP-A nucleosomes.

## Interactions between CENP-N α6 and DNA of a neighboring nucleosome promote nucleosomes stacking *in vitro*

How does CENP-N interact with the second nucleosome? Our structures reveal an additional, previously unidentified interface between CENP-N and nucleosomal DNA, consisting of a series of positively charged amino acids (K102, K105, K109, K110, R114, and K117) that are all located on the same face of the α6 helix of CENP-N, and on the opposite side of the main CENP-A decoding interface on CENP-N. These allow α6 to dock onto ‘super helix location’ (SHL) 4-5 of the second nucleosome in the stack (fig. 2A). Consistent with the electrostatic nature of this interface, nucleosome stack formation is strongly affected by ionic strength (fig. S5). At 200 mM salt, CENP-N is still able to interact with the CENP-A nucleosome but no stack formation is observed. Point mutation of individual side chains (K102A or R114A) did not affect the specific interaction between CENP-N and CENP-A mono-nucleosomes (fig. S6A) but resulted in reduced levels of nucleosome stacking (fig. 2B). This is consistent with our MD simulations which demonstrate that the α6 helix (in particular the amino acids listed above) form strong contacts with the neighboring DNA (fig. S7). The charged face of the α6 helix is not conserved in CENP-N from fungi with point centromeres, and this may reflect the dispensibility of an additional bridging interface between nucleosomes in a point centromere compared to a regional centromere (*24*).

**Fig. 2.**
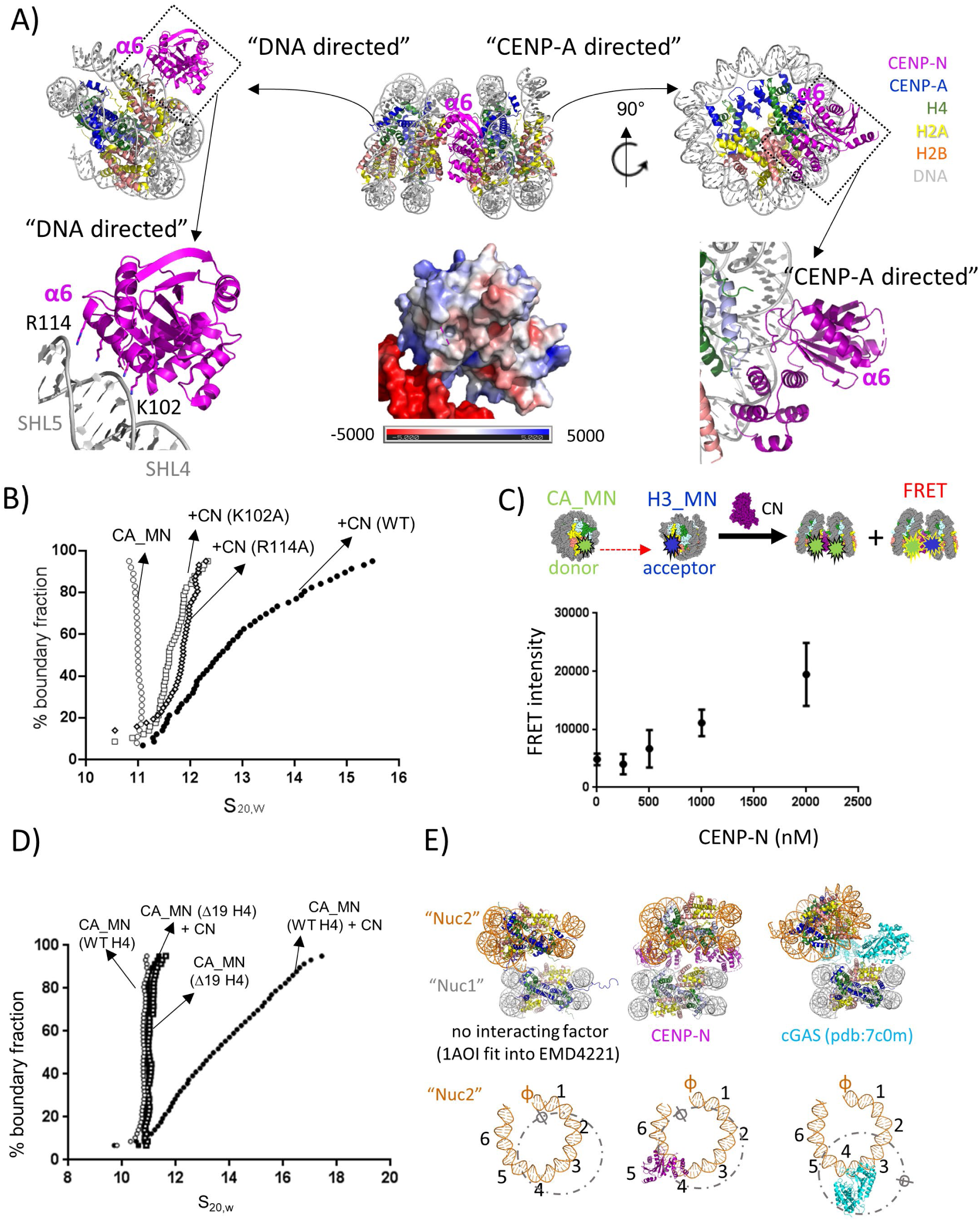
Structural basis for CENP-N dependent nucleosome-nucleosome interaction. **A)** The CENP-N α6 helix interacts with nucleosomal DNA of a second nucleosome without contacting histones (‘nonspecific nucleosome’). Upper panel: overview of interactions. Lower left panel highlights the interface between CENP-N α6 and DNA. Lower middle panel shows a surface charge representation (unit : kT e^−1^) in the same orientation. Lower right panel highlights the specific interaction on the other side of CENP-N with the CENP-A RG loop (*13*). **B)** Single mutations on CENP-N α6 affect nucleosome-nucleosome interactions, as shown by AUC. **C)** FRET analysis of the interaction between CENP-A mono-nucleosome (CA_MN) and H3 mono-nucleosome (H3_MN) in the absence or presence of CENP-N. Donor is CENP-A mono-nucleosomes containing Alexa 488 labeled H2B; Acceptor is a H3 mono-nucleosome containing Atto N 647 labeled H2B. 250 nM donor and acceptor nucleosome; FRET intensity in dependence of [CENP-N]. Error bars from five independent measurements of two biological repeats. **D)** The H4 N-terminal tail is essential for CENP-N mediated nucleosome stacking (van Holde Weischet plots of sedimentation). Δ19 indicates H4 tail deletion (amino acids 1-18). **E)** comparison of different modes of nucleosome stacking. Upper panel shows models for stacked mono-nucleosomes. Lower panel indicates the Super Helical Locations (SHL) and the nucleosomal dyad axis (SHL 0; ɸ) of nuc2, with the orientation of the nuc1 shown in a dotted circle and depicts the nonspecific interaction with CENP-N and cGAS, respectively. Only half of the nucleosomal DNA is shown for clarity, and the dyad axis of the ‘specific nucleosome’ is indicated by ɸ.

Since one CENP-N is sufficient to mediate nucleosome stack formation, and since the second nucleosome interacts with CENP-N via its DNA, it could, in theory, also promote stacking between CENP-A and H3 nucleosomes. To test this, we performed FRET experiments with CENP-A- and H3-nucleosomes labeled with fluorescence donor and acceptor, respectively. Weaker but significant FRET signal was observed between the CENP-A nucleosome and H3-nucleosome with increasing CENP-N concentrations, confirming our prediction (fig. 2C).

Histone tails, especially the H4 N-terminal tail, contribute to chromatin compaction (e.g. (*25–28*)). Because CENP-N appears to redirect the H4 tail (*13*), we tested by AUC whether the H4 tail contributes to CENP-N mediated nucleosome stacking. We prepared CENP-A-containing nucleosomes in which the H4 N-terminal tail was deleted (Δ19H4). No CENP-N dependent oligomerization was observed for these nucleosomes (fig. 2D), even though they bind to CENP-N as well as CENP-A nucleosomes containing full length H4 tails (fig. S6B).

Nucleosome-nucleosome interactions have been observed previously. For example, major-type nucleosomes form several types of dinucleosomes on cryoEM-grids, in absence of any interacting protein ((*29*) and fig. 2E), likely mediated through histone tails. Recent cryoEM structures of cGAS (a protein that senses the presence of cytoplasmic DNA during the innate immunity response) in complex with nucleosomes show that it bridges two mono-nucleosomes. cGAS binds to one nucleosome by interacting with the surface of histones H2A-H2B and nearby DNA, while a positively charged α-helix interacts with DNA of the second nucleosome (fig. 2E) (*30–32*). A caveat here is that the existence of these nucleosome stacks was not verified in solution. Of note, the relative orientation of the nucleosomes is quite different in the three arrangements (second nucleosome indicated by dashed circles in figure 2E), enforcing the concept gained from nucleosome crystallography that there are many ways to pack nucleosomes in an energetically favorable way (e.g. (*33*)).

## CENP-N folds and twists CENP-A containing chromatin arrays

The interactions between mono-nucleosomes observed *in vitro* might reflect how nucleosomes form long-range interactions *in vivo* without constraints from connecting DNA. We next asked whether CENP-N promotes the short-range interactions required to form chromatin fibers from a linear nucleosomal array. CENP-N^1-289^ was mixed with CENP-A-containing arrays assembled onto 12 tandem repeats (12mer) of 207 or 167 bp 601 DNA, at a ratio of five CENP-N^1-289^ per nucleosome to reach saturation. CryoEM images showed that both chromatin arrays fold into twisted zig-zag chromatin fibers (fig. S8A, B). This type of folding is usually only observed in the presence of divalent cations or upon addition of linker histone to canonical H3 arrays (*26, 34*). While the longer linker segments in 12-207 nucleosomal arrays introduced too much variability to allow structure determination, we were able to determine the ∼12.7 Å structure of the more constrained 167-12-mer CENP-A array in complex with CENP-N (fig. 3A). Of note, the average linker length at the centromere is ∼ 25 bp (*35*), close to the linker length of ∼20 bp used here. Eight nucleosomes, each bound by two CENP-N molecules, were observed in the density map; the two terminal nucleosomes on either end were too flexible to be described with any certainty.

**Fig. 3.**
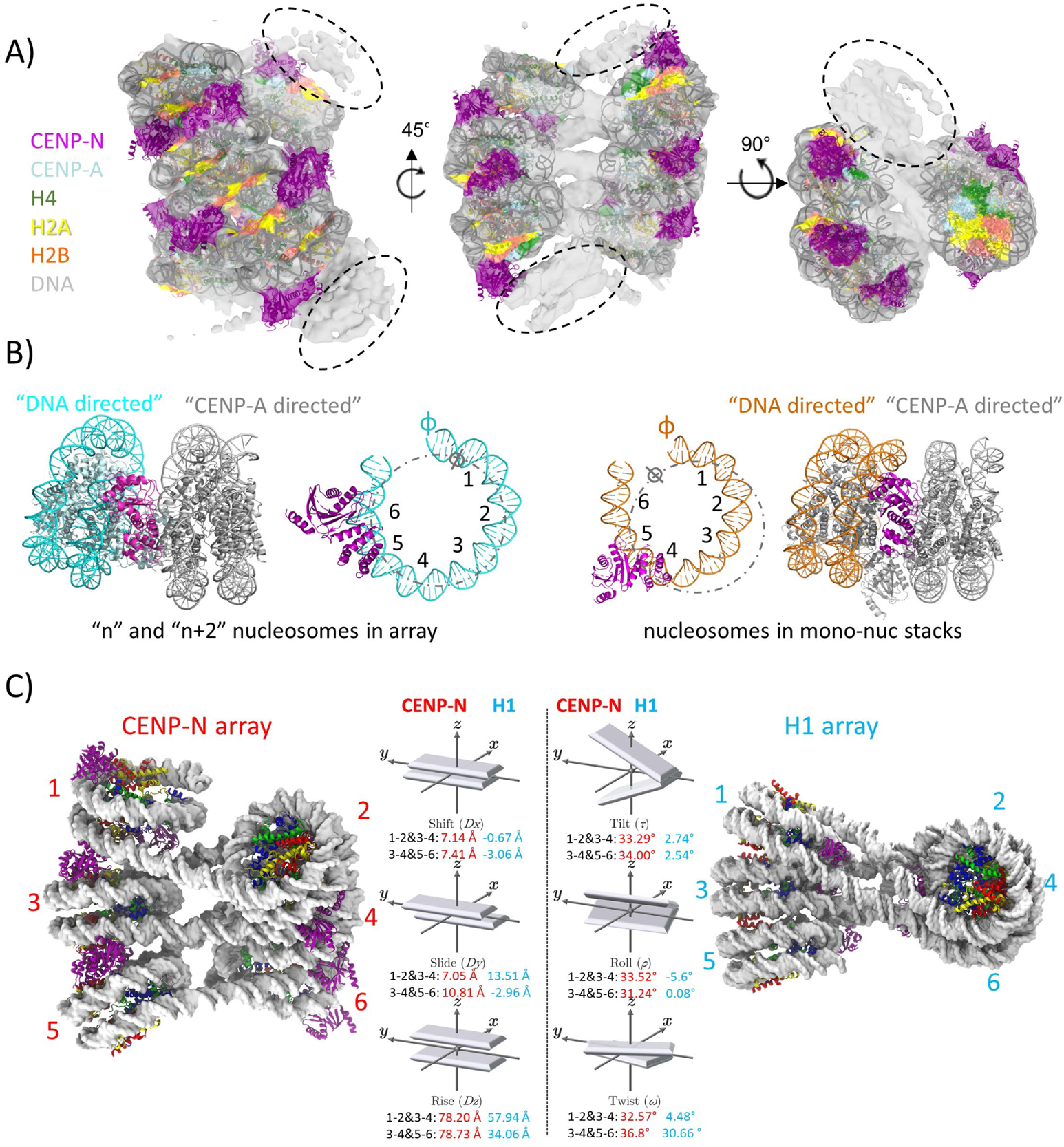
CENP-N folds CENP-A chromatin into a regular fiber. **A)** A nucleosomal array consisting of 6 CENP-A nucleosomes with CENP-N was fit into the density map. Dashed circles indicate weak densities representing more disordered nucleosomes at both ends. **B)** CENP-N interacts with nucleosomal DNA of a neighboring nucleosome (CENP-N α6 binds between SHL6 and 7 (left panel)). This is in contrast to mono-nucleosome stacks, where CENP-N α6 binds between SHL4 to 5 (right panel). **C)** the relative arrangement of nucleosomes in the arrays was expressed in analogy to the arrangement of DNA bases in a double helix. Parameters for CENP-N fibers (left, EMDxxx) and H1 fibers (*26*) are shown. The vectors in the middle show the parameters of the analysis (in red for CENP-A arrays, and cyan in H1 arrays).

The arrangement of nucleosomes takes the form of a two-start twisted double helix with two CENP-N bridging n and n+2 nucleosomes. CENP-N is in its previously described location on the CENP-A nucleosome and interacts with the DNA of the n+2 nucleosome, but binds SHL 6 and 7 of the n+2 nucleosome rather than SHL 4 and 5 as observed in mono-nucleosome stacks (fig. 3B). This results in a different alignment of the n, n+2 nucleosome stack compared to that formed from mono-nucleosomes, and provides evidence for the plasticity of the interaction between α6 and nucleosomal DNA.

Linker histone H1 (which binds to the nucleosomal dyad and linker DNA (*36*)) stabilizes compact chromatin states (*37*). Although the manner in which CENP-N and H1 interact with nucleosomes are completely different, they both promote chromatin fibers with superficially similar two-start zig-zag architectures held together by the stacking of n and n+2 nucleosomes (fig. 3C). In molecular dynamics simulation, chromatin converts to a parallel “ladder like” structure when H1 is removed from the simulation (*38*). We observed a similar structural change by cryoEM when CENP-N was lost in buffer containing 200 mM NaCl during overnight sucrose gradient centrifugation (fig. S9). Interestingly, a similar “ladder like” structure on CENP-A chromatin array was also observed in the presence of divalent cations (*39*). As such, the structural transition caused by CENP-N in CENP-A arrays is very similar to that caused by H1 on canonical chromatin. Intriguingly, H1 is unable to bind to CENP-A nucleosomes *in vitro* and *in vivo* (*9, 10*), and we speculate that CENP-N might take over the role of H1 in closely packing CENP-A nucleosomes with surrounding nucleosomes.

The CENP-N / CENP-A chromatin fiber exhibits features that distinguish it from the H1-induced fiber. A larger distance and angle between the n and n+2 nucleosome is required to accommodate CENP-N. This leads to a steeper twist of the fiber (fig. 3C). Additionally, the chromatin fiber formed with H1 exhibits a discrete “tetra-nucleosomal structural unit”, a repeat of four nucleosomes (*26*), while the organization of CENP-A chromatin fibers with CENP-N is continuous. Of note, the packing of n and n+2 CENP-A nucleosomes in the presence of CENP-N also differs from the nucleosome interactions observed in the crystal structure of a canonical tetranucleosome stack (*34*) (fig. S10).

CENP-A nucleosomes are characterized by less-tightly bound DNA ends, which affects the geometry of CENP-A containing chromatin (*8*). In the presence of CENP-N, all CENP-A nucleosomes (both in mono-nucleosome stacks and in folded chromatin arrays) exhibit tightly bound DNA ends, similar to what is observed for canonical nucleosomes (this study and (*13*)), and in this CENP-N also functionally resembles linker histone H1. Overall, our data suggests that CENP-N, as one of the key proteins of the inner kinetochore, stabilizes, organizes and compacts centromeric chromatin in a way that depends on its specific interaction with CENP-A nucleosomes and on its “DNA-directed” interactions with a neighboring nucleosome.

## CENP-N promotes the compaction of centromeric chromatin *in vivo*

To explore the role of CENP-N in the compaction of centromeric chromatin *in vivo*, we used sucrose gradient ultracentrifugation to fractionate and separate mechanically sheared cross-linked chromatin isolated from cells. As shown previously, chromatin domains with higher levels of compaction (e.g. heterochromatin) resist sonication more effectively than the more open euchromatin, and thus sucrose gradient ultracentrifugation enables separation of these different chromatin states. The sonication-resistant, more compact chromatin migrates faster and sediments in fractions of high sucrose density (*40, 41*), whereas more open chromatin migrates slower and fractionates at lower sucrose density.

Here, we used a 5-40% sucrose gradient, and identified the fractions containing centromeric chromatin with antibodies against CENP-A, CENP-N, and CENP-C in Western blots. We observed that centromeric chromatin (anti hCENP-A antibody signal) sediments in high-density sucrose fractions (e.g. 12-20) (fig. 4A, shown in gray). These same fractions also contain highly compacted heterochromatin as they also stain with antibodies against H3K9me3, a marker of constitutive heterochromatin (fig. S11A) (*42–44*). This suggests that centromeric chromatin indeed resists sonication just like heterochromatin, reflecting a high level of compaction.

**Fig. 4.**
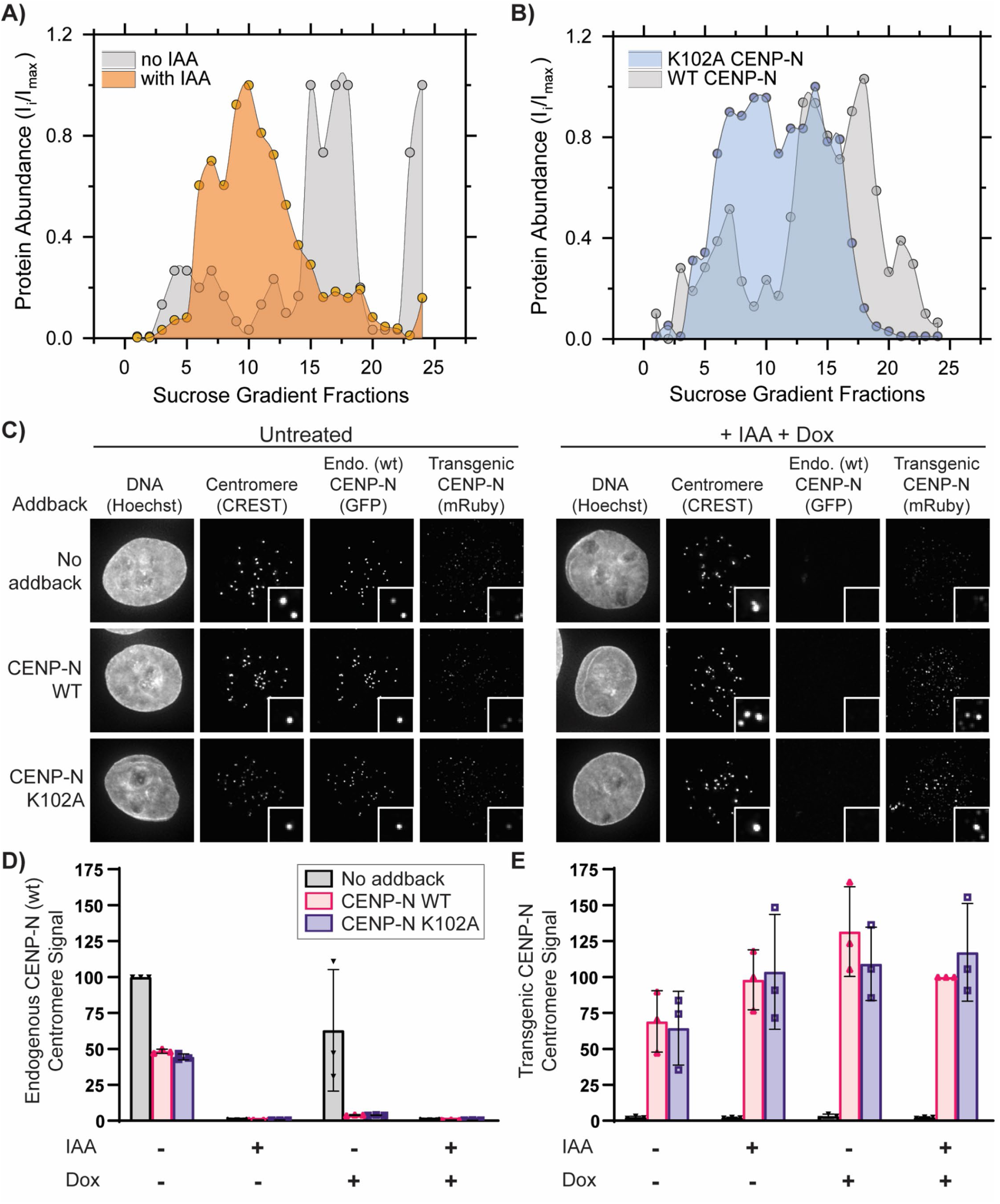
CENP-N promotes the compaction of centromeric chromatin *in vivo.* **A**) CENP-A distribution in sucrose gradient in the presence of WT-CENP-N (gray) and after IAA-induced degradation of CENP-N (orange). **B**) Comparision of CENP-A distribution in the presence of transiently exressed CENP-N variants, WT CENP-N (gray) and K102A CENP-N (lilac). Dots in **A & B** represent a mean of two biological replicates. Solid lines represent interpolation between experimental data points. **C)** Representative images of nuclei (Hoechst stain) showing endogenous wildtype CENP-N (GFP) or transgenic CENP-N variant (anti-Ruby antibody) localization at centromeres (CREST signal). Insets show magnified images of example centromeres (CREST foci). Cells were treated for 24 hours with doxycycline (DOX) to induce expression of mRuby2-3xFLAG-tagged CENP-N (WT or mutant) and/or with Indole Acetic Acid (IAA) to deplete endogenous Auxin Inducible Degron (AID) tagged CENP-N-GFP as indicated. **D** & **E)** Normalized centromeric fluorescence signal corresponding to endogenous WT CENP-N (GFP) [D] or transgenic WT or CENP-N variant (anti-mRuby antibody) [E]. Each dot represents median centromeric CENP-N signal from >100 cells from one biological replicate. Line and error bars are mean of 3 biological replicates and standard deviation respectively.

To assess the role of CENP-N in compacting CENP-A nucleosomes *in vivo*, we endogenously tagged both alleles of CENP-N with the auxin-inducible degron (AID) tag in cells expressing the F-box protein, Tir1. This AID system enables targeting CENP-N for degradation upon the addition of auxin (*15*). Degradation of endogenous CENP-N by auxin addition causes depletion of CENP-N from centromeres (fig. 4C, D) and a reduction in long-term cell viability (fig. S12) (14). Transient degradation of CENP-N for 24 hours caused significant changes in the migration of centromeric chromatin in sucrose gradients, assayed by CENP-A distribution. Most CENP-A chromatin from these cell lines now migrates with lower-density sucrose gradients (fractions 5-15, fig. 4A, in orange), indicating that the loss of CENP-N renders centromeric chromatin more accessible to shearing by sonication. We tested if transgenic expression of CENP-N rescues the effects of endogenous CENP-N degradation on CENP-A chromatin migration by introducing into AID-tagged CENP-N cells mRuby2-3xFLAG-tagged full length CENP-N as a transgene (transgene WT-CENP-N) under doxycycline induction. Upon doxycycline addition, we observed localization of transgenic CENP-N at centromeres by immunofluorescence using anti-Ruby antibody in cells depleted of endogenous CENP-N (fig. 4C-E). Introduction of CENP-N WT transgene in CENP-N AID cells significantly reduced the centromeric localization of endogenous CENP-N, suggesting that transgenic CENP-N WT competes with endogenous CENP-N for centromere localization (fig. 4D,E). Furthermore, transgenic CENP-N WT was able to rescue loss of cell viability in CENP-N AID cells (fig. S12), even in the absence of doxycycline, suggesting leaky background expression of transgenic CENP-N. Indeed, we observed significant anti-Ruby signal at centromeres in transgenic CENP-N WT addback cells even in the absence of doxycycline (fig. 4E). Complementation of AID-CENP-N loss through WT CENP-N expression rescues the migration of CENP-A nucleosomes in sucrose gradients to what is observed in unmanipulated cells (fig. S11B). The distribution of transgenic CENP-N in the gradient overlapped with the distribution of the CENP-A nucleosomes indicating that transgenic CENP-N associates with CENP-A chromatin (fig. S11C). Overall, these results demonstrate a role for CENP-N in promoting the compaction of centromeric chromatin in the cell.

We next tested the contribution of the α6 helix of CENP-N to the compaction of centromeric chromatin by complementing AID-CENP-N degradation with transgenic expression of mRuby2-3xFLAG-tagged K102A mutant of CENP-N. Upon doxycylcine addition, CENP-N K102A localized to centromeres in cells depleted of endogenous CENP-N (fig. 4C-E) indicating that it maintains its ability to target to CENP-A chromatin in cells. Moreover, similar to what was observed for transgenic expression of WT CENP-N, transgenic expression of CENP-N K102A restored long-term viability in cells depleted of endogenous CENP-N (fig. S12). This is in contrast to mutations in the CENP-N/CENP-A histone interface which perturb CENP-N localization and cell viability (*13, 45*). However, complementation of AID-CENP-N depletion with the CENP-N α6 mutant caused centromeric chromatin to become less resistant to shearing, similar to what was observed upon degradation of endogenous CENP-N. Most CENP-A containing chromatin was localized in fractions 5-15 in the presence of the CENP-N α6 mutants as compared to fractions 12-20 in the presence of WT CENP-N (fig. 4A, B). We confirmed that transgenic CENP-N mutant proteins were indeed expressed and that their distribution within the sucrose gradient overlaps with the corresponding CENP-A distribution (fig. S11D, E).

Like CENP-N, CENP-C also directly binds to CENP-A nucleosomes (*45*). We found that the distribution of CENP-C in sucrose gradients is different in cell lines expressing CENP-N mutants compared to cell lines with WT CENP-N (fig. S11E, G). CENP-C migrated with low-density sucrose fractions that contain CENP-A and K102A CENP-N, but we also detected CENP-C in high-density sucrose fractions which were depleted in CENP-N but not in CENP-A (fig. S11G). These results suggest that there is a population of CENP-A nucleosomes not bound by CENP-N that sediments with compacted centromeric chromatin (e.g. high-density sucrose fractions, fig. S11D, G, fractions 15-20) and that this compaction state corresponds to the presence of the CENP-C protein. Altogether, our *in vivo* studies validate the *in vitro* results to demonstrate that CENP-N plays an important role in compaction of CENP-A nucleosomes through its α-6 helix.

## Discussion

The specialized chromatin structure at the centromere, which is necessary for the assembly of the kinetochore, is initiated by two proteins (CENP-N and CENP-C) that specifically interact with CENP-A containing nucleosomes. These are found only at centromeres, where they are interspersed with H3-nucleosomes along the centromeric DNA, but clustered in 3D. CENP-N promotes the stacking of nucleosomes *in vitro* and *in vivo* through a previously undescribed DNA interaction interface, and this has important implications for our understanding of higher order structure at the centromere and for the transmission of force on chromosomes exerted by mitotic spindle fibers. Our finding that the H4 tail contributes to nucleosome-nucleosome interactions is underscored by the fact that histone H4 at the centromere is not acetylated in its tail regions (*46*). This is notable because previous data showed that acetylation of H4K16 precludes the formation of higher order structure *in vitro* and *in vivo* (*47*). The interactions between CENP-N and the CENP-A nucleosome that we observe here and in (*13*) are different from those previously described for the yeast CCAN complex (*48*). It will be interesting to determine whether this reflects an evolutionary adaptation for stabilizing point versus regional centromeres.

While CENP-N is specific for CENP-A nucleosomes (*49*), the interaction with the neighboring nucleosome appears to be promiscuous with respect to histone content and DNA sequence / location. Thus CENP-N can promote the close packing of CENP-A nucleosome and its varied surrounding nucleosomes even including sub-nucleosomes (hexasomes or tetrasomes). CENP-N promotes the formation of stacks from mono-nucleosomes (reflecting the interaction of unconnected nucleosomes from different regions of the genome), as well as nucleosomal arrays, where it leads to the formation of a zig-zag two-start helix with a topology that is distinct from the canonical nucleosome fiber formed by linker histone H1. This is significant as H1 is indeed unable to bind to CENP-A nucleosomes. It is therefore possible that CENP-N acts as a centromere-specific ‘linker histone’ to promote the formation of centromere-specific chromatin higher order structure, which in turn serves as an interaction platform for a plethora of additional centromere-specific proteins.

Chromatin higher order structures are heterogeneous *in vivo* and can be influenced by a variety of factors. Linker histone H1 compacts chromatin by organizing the extranucleosomal linker DNA. Heterochromatin protein-1 (HP1) promotes heterochromatin formation through reading the histone modification H3K9me3 as well as self-dimerization (*50*). Here we reported yet another chromatin compaction mechanism where CENP-N specifically reads the histone variant content on one nucleosome while interacting with the DNA on a neighboring nucleosome, to potentially form unique chromatin structures at the centromere.

**Table 1.**
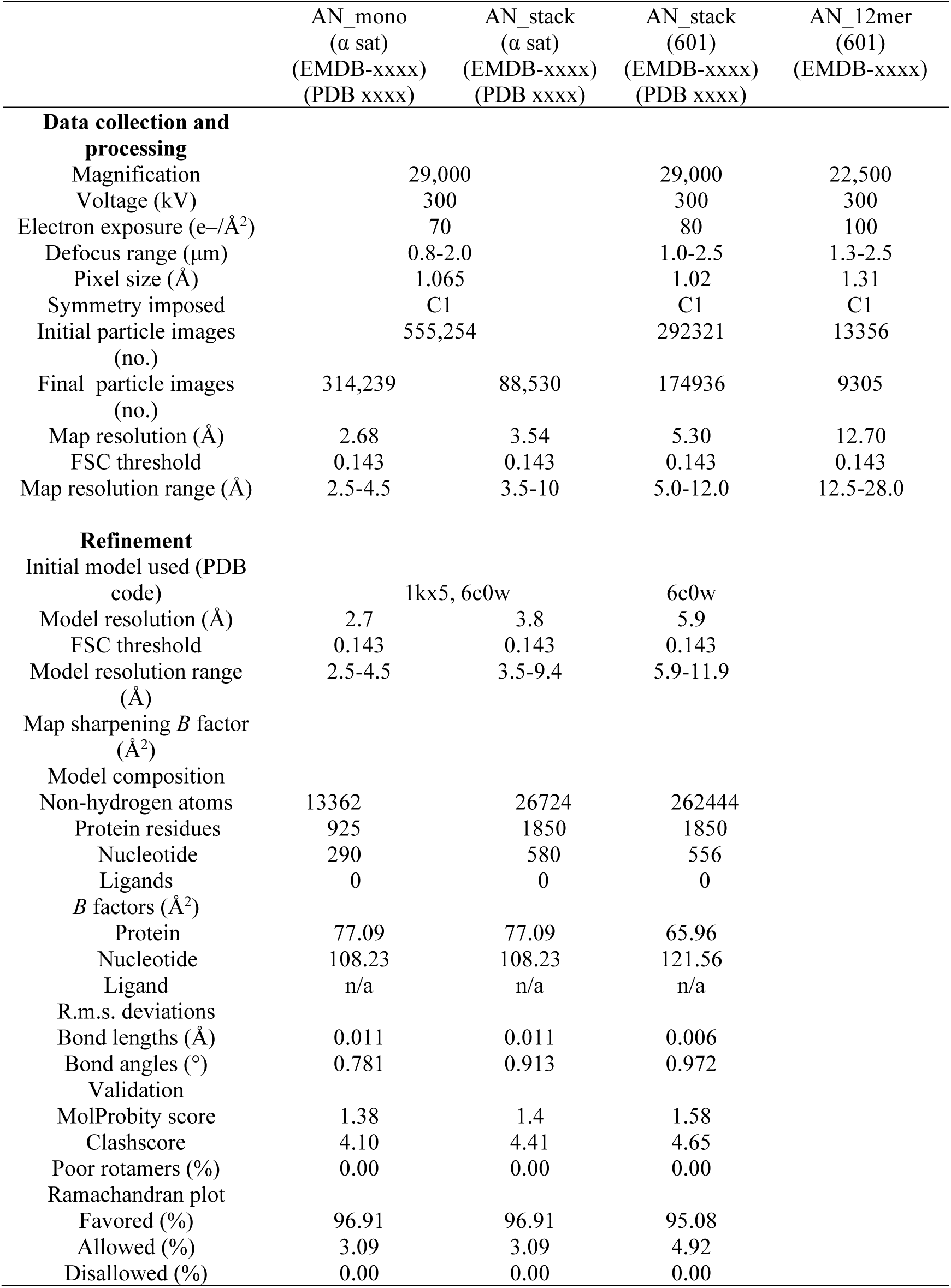
Cryo-EM data collection, refinement and validation statistics.

## Acknowledgements

We thank Andrea Musacchio for the kind gift of CENP-N/CENP-L protein, and Alison White for preparation of DNA, and Yang Liu for CENP-N purification. CryoEM data were collected at the CryoEM Facility at the HHMI Janelia Research Campus (Doreen Matthies and Zhiheng Yu) and at the CU Boulder BioKEM facility (Charles Moe).

## Funding

Howard Hughes Medical Institute (KZ, KL)

National Institutes of Health Grant R01GM74728 (AFS)

National Institute of Health Grant R35GM119647 (JW)

National Science Foundation Grant 1552743 (JW)

## Author contributions

Conceptualization: KZ, KL, AS, MG, KS

Methodology: KZ, KL, MG, KS, DW, JW, AS, GE, DK

Investigation: KZ, KL

Visualization: KZ, KL, MG, KS, DW, JW, AS

Funding acquisition: KL. JW, AS

Project administration: KL, AS

Supervision: KL. JW, AS

Writing – original draft: KZ, MG, KS, KL

Writing – review & editing: KZ, KL, MG, KS, DW, JW, AS, GE

## Competing interests

Authors declare that they have no competing interests.

## Data and materials availability

All data, code, and materials will be made freely available. CryoEM maps will be deposited at EMDB, and pdb files of mono- and dinucleosome structures (with appropriate validation) will be deposited in the pdb. Simulation trajectories are available on request.

## Supplementary Materials

### Materials and Methods

#### Protein expression and purification

CENP-N^1-289^ was cloned into pACEBac1 with a (his)6 tag at its C-terminus. It was expressed in Sf21 insect cells (Invitrogen, Thermo Fisher Scientific, USA), and purified through affinity tag (nickel beads), followed by size-exclusion column (S200) as described (*13*). Full length CENP-N:CENPL was a gift from Andrea Musacchio (MPI Dortmund). Histones H3, H4, H2A and H2B were purchased from ‘The Histone Source’ at Colorado State University. (H3-H4)_2_ and H2A-H2B were refolded as described (*22*). H2B T112C was labeled by Alexa 488 or Atto N 647 during the refolding procedure as previously described (*22*). CENP-A and H4 were co-expressed from a bi-cistronic plasmid in BL21(DE3)pLysS cells. Two-step purification over hydroxyapatite and Hitrap SP FF yielded pure, folded (CENP-A:H4)_2_ (*51*).

#### Nucleosome reconstitution

Nucleosomes were reconstituted using the salt dialysis method as described (*22*). DNA, H2A-H2B dimer and (CENP-A-H4)_2_ or (H3-H4)_2_ tetramer were mixed at molar ratios of 1.0:2.4:1.2 in reconstitution buffer A (Tris-HCl pH 7.5, 2 M NaCl, 1 mM DTT, and 1 mM EDTA) and dialyzed against buffer A. Using a peristaltic pump (Econo pump, Bio-Rad, Richmond, CA), buffer B (Tris-HCl pH 7.5, 50mM NaCl, 1 mM DTT, and 1 mM EDTA) was exchanged with buffer A at the rate of 1.5 ml/minute overnight (>16 hours). The nucleosomes were then dialyzed against buffer B for 2 hours. Nucleosome quality was monitored by native PAGE.

#### Analytical Ultracentrifugation (AUC)

Sedimentation Velocity AUC (SV-AUC) was performed with absorbance optics (λ = 260 nm) to evaluate the homogeneity, size and shape of nucleosome and its complex with CENP-N. The nucleosome concentration in SV-AUC was fixed at 250 nM (OD260∼0.5). CENP-N was added at a ratio of 3:1 or higher, as indicated. The buffer contained 20 mM Tris-Cl (pH7.5), 1 mM EDTA, 1 mM DTT and NaCl at different concentration (50 mM or 200 mM). For mono-nucleosome samples, the centrifugation speed was 30-35,000 rpm at 20°C in a Beckman XL-A ultracentrifuge (An60Ti rotor).

UltraScan III version 4.0 (*52*) was used for the data analysis. 2-dimensional Spectrum Analysis (2DSA; (*53*)) subtracted time-invariant and radial-invariant noise contributions from the experimental AUC data. Enhanced van Holde-Weischet Analysis method was then performed to obtain Integral sedimentation coefficient distributions (G(s)) from noise-corrected experimental data. The AUC figures were Enhanced van Holde-Weischet plots (*54*).

#### Electrophoretic Mobility Shift Assay (EMSA)

5% native PAGE was used to monitor the nucleosome quality as well as binding events between CENP-N and nucleosomes (*15*). The buffer for binding assays contained 20 mM Tris-Cl (pH7.5), 1 mM EDTA, 1 mM DTT and 50 mM NaCl.

#### Forster Resonance Energy Transfer (FRET)

CENP-A or H3 nucleosomes were reconstituted with Alexa 488 or Atto N 647 labeled H2B (T112C). Alexa 488 served as the donor while Atto N 647 as the acceptor. Equal amount (250 nm) of donor nucleosome and acceptor nucleosome were mixed in the absence or presence of CENP-N (molar ratio to CENP-A nucleosome at 2:1 and 4:1). FRET signal was obtained with the excitation wavelength at 482(+/-16) nm and emission wavelength at 675(+/-50) nm. The FRET value were then corrected as described (*55*). The signal from the emissions of donor and acceptor in FRET channel were removed from the raw FRET value.

#### CryoEM sample and grid preparation

CENP-N and CENP-A mono-nucleosome were mixed at a 5:1 ratio in buffer containing 20 mM Tris-Cl (pH7.5), 1 mM EDTA, 1 mM DTT and 50 mM NaCl. The complex was concentrated to 1-2 µM before grid preparation. For grid plunge freezing, 4 µl sample was loaded on a holy carbon grid (Quantifoil 1.2/1.3 Au) and flash frozen using a Vitrobot Mark IV (ThermoFisher Scientific). Blotting time was 4 s with >90% humidity.

CENP-N and CENP-A nucleosome arrays (‘601’ sequence, 167bpx12mer) were mixed at a 5:1 (nucleosome) ratio in buffer containing 20 mM Tris-Cl (pH7.5), 1 mM EDTA, 1 mM DTT and 300 mM NaCl. The buffer for the mixture was gradually exchanged/dialyzed against buffer the same buffer containing 50 mM NaCl (overnight at 4°C). The sample was then concentrated to 0.1-0.2 mg/ml before grid preparation. For grid plunge freezing, 4 µl sample was loaded on a holy carbon grid (C-Flat 2/2 Au) and flash frozen using a Vitrobot Mark IV (Thermo Scientific). Blotting time was 4 s with >90% humidity.

The structure of a crosslinked CENP-A nucleosomal array (‘601’ sequence, 167×12mer) was an accidental product of an attempt to GRaFix (gradient fixation by glutaraldehyde) CENP-N and CENP-A nucleosome array in buffer containing 20 mM HEPES (pH7.5), 1 mM EDTA, 1 mM DTT and 200 mM NaCl overnight, which caused CENP-N to dissociate. Fractions were combined to be quenched by dialyzing against the buffer containing 20 mM Tris-Cl (pH7.5), 1 mM EDTA, 1 mM DTT and 50 mM NaCl for >2 hours. The sample was concentrated to 0.1-0.2 mg/ml before grid preparation. For grid plunge freezing, 4 µl sample was loaded on a holey carbon grid (C-Flat 2/2 Au) and flash frozen using a Vitrobot Mark IV (Thermo Scientific). Blotting time was 4 s with >90% humidity.

#### CryoEM images acquisition and analysis

CENP-N in complex with CENP-A mono-nucleosome (alpha satellite DNA) was imaged at nominal magnification of 29000x on a FEI Titan Krios (300 kV), equipped with a Gatan K3 direct detector. Pixel size was 0.8211 Å. The movies were captured in counting mode with electron dose rate at 10 electrons per pixel per second for 7.2 s and 0.2 s per frame. Defocus range was −1.0 to −2.5 µm. The raw movies were motion corrected, followed with Constant transfer function (CTF) estimation in cryoSPARC2 (*56*). Nucleosome like particles were manually picked to generate the templates for “template picking”. 555,254 particles were picked by “template picking”. The box size of particle extraction was set at 400 pixels. One round of 2D classification was performed to remove the low resolution or non-nucleosome particles. Ab initio classification was used to generate the initial maps (3 classes). One class (class 1) was mainly at mono-nucleosome level. The other two classes (class 2&3) were both di-nucleosome organization. Class1 was continually processed by 2D classification as well as heterogeneous refinement to remove low resolution particles. Non-uniform refinement (*57*) was performed to obtain the highest resolution map for class1. For class2 and class3, they looked very similar to each other. Therefore, the particles from these two classes were combined to be processed by 2D classification and heterogeneous refinement to remove low resolution or unaligned particles. The model was finally refined by non-uniform refinement (*57*). 3DFSC was used to evaluate and reflect the directional resolution (*58*). The map for complex in di-nucleosome form showed very heterogeneous resolutions in different direction due to particle orientation problems.

CENP-N in complex with CENP-A mono-nucleosome (601 DNA) was imaged at nominal magnification of 29000x on a FEI Titan Krios (300 kV), equipped with a Gatan K2 Summit direct detector. Pixel size was 1.02 Å. The movies were captured in super resolution mode with electron dose rate at 10 electrons per pixel per second for 8 s and 0.2 s per frame. Defocus range was −1.0 to −2.5 µm. The raw movies were motion corrected by Motioncor2 (*59*), followed with Constant transfer function (CTF) estimation through GCTF (*60*). To classify nucleosome pairs in the stack, the size of pixel box was enlarged to 256 which was enough to include two nucleosomes. 2D classifications were performed in cryoSPARC2.0 to select the 2D classes representing nucleosome stacks. Ab initio reconstruction was used to generate the initial model, followed by two rounds of heterogenous refinement (two classes). Both classes were refined by non-uniform refinement (*57*). 3DFSC analysis was performed to evaluate the directional resolution (*58*).

Chromatin fiber with CENP-N was imaged at nominal magnification 22500x by a FEI Titan Krios (300 kV),equipped with a Gatan K2 Summit direct detector. The images were captured in super resolution mode (pixel size 0.655 Å) with electron dose rate at 10 electrons per pixel per second for 10 s. Defocus range was −1.3 to −2.5 µm. The raw movies were motion corrected, followed with Constant transfer function (CTF) estimation in cryoSPARC2 (*56*). Cryolo was used for particle picking (*61*). Extracted particles were imported into cryoSPARC2. 2D classifications were performed to remove the junk particles. Ab-Initio reconstruction (2 classes) was performed to generate the initial map and remove the junk particles. Non-uniform refinement was used to do the final refinement (*57*). 3DFSC was used to evaluate the global resolution (*58*).

Crosslinked chromatin fiber was imaged at nominal magnification of 22,500 on a FEI Titan Krios (300 Kv), equipped with a Gatan K3 direct detector. The images were captured in counting mode (pixel size 1.064) with electron dose rate at 10 electrons per pixel per second for 7.8 s (65 frames with 0.12 s per frame). Defocus range was −0.9 to −2.1 µm. The raw movies were imported and processed by motion correction and CTF estimation in cryoSPARC2 (*56*). Folded chromatin particles were manually picked to generate the templates for “template picking”. 2D classifications were performed to remove the junk particles. Ab-Initio reconstruction (2 classes) was performed to generate the initial map and remove the junk particles. Non-uniform refinement was used to do the final refinement (*57*).

#### Modeling of the structures

The structure of CENP-N in complex with a mono-nucleosome (α-satellite DNA) was generated by combining CENP-N and histones from pdb:6c0w (*13*) and the α-satellite DNA from pdb: 1kx5(*62*). DNA was shifted manually by 1 bp as indicated in supplementary text. MDFF was performed to refine the model (*63*). The structure model for CENP-N in complex with di-nucleosome was initially generated by docking the model of CENP-N in complex with mono-nucleosome twice in the map. Although the map shows high heterogeneity on the resolution in different direction, it did not affect unambiguous docking of the model.

The structure model of CENP-A chromatin fiber with CENP-N was generated by placing 6 copies of pdb:6c0w (6 times) into the map.

#### Generation of cell lines

CRISPR-Cas9 based genome engineering was used to tag CENP-N with AID-sfGFP at the C-terminus as descripted previously (*30*). Briefly, native CENP-N locus in osTir1 Flp-In TRex-DLD1 cell line was tagged with AID-sfGFP in the C-terminus. Transgenic CENP-N constructs (WT full length CENP-N and CENP-N K102A mutant) with C-terminus with tandem mRuby2 and 3xFlag were induced using FRT/Flp-mediated recombination of pcDNA5/FRT/TO-vectorand selected with 100 µg/ml Hygromycin B. WT full length CENP-N was expressed from ASP 3329 (*15*). CENP-N K102A mutation in CENP-N was introduced in ASP 3329 using Gibson Assembly (CENP-N K102A Plasmid ASP 4243; forward Gibson oligo: GGACGTGGATCTGTTCGACATGGCGCAGTTCAAGAACAGCTTTAAAAAG reverse Gibson oligo: cgaggacgtggatctgttcgacatgGCGcagttcaa-gaacagctttaaaaagatc) and confirmed by sequencing. Full plasmid sequences are available from the Stanford Digital Repository (https://purl.stanford.edu/gz478gq3828).

#### Cell culture

All cell lines were grown in RPMI-1640 medium supplemented with 10 % fetal bovine serum (FBS), 100 U/mL penicillin/0.1 mg/mL streptomycin, 2 g/L sodium bicarbonate, and 2 µg/mL puromycin. Cells were tested for mycoplasma contamination at the start of cell line generation by PCR and conditions were monitored by the absence of cytoplasmic DNA staining by microscopy. Degradation of the endogenous AID-tagged CENP-N was carried out by adding 1 mM 3-Indoleacetic acid (IAA) (Sigma, # I2886), followed by 1 h incubation. Expression of transgenic CENP-N mutants was induced using 20 ng/mL doxycycline (Sigma, #D9891) accompanied by degradation of the endogenous CENP-N using 1 mM IAA, followed by 24 h incubation. All drugs were added directly to the media.

#### Preparation of crosslinked chromatin lysates

Cell were grown to 80 % confluence in 150-mm dishes (10 plates per chromatin batch, 70 – 80 · 106 cells). Cells were trypsinized, twice pelleted at 1000 g for 5 min and washed in PBS buffer. Crosslinking was carried out by addition of formaldehyde (16 % formaldehyde (w/v), methanol-free, Thermo Scientific, #28908) to 10 mL of PBS buffer for 1 % formaldehyde final concentration, rocking for 10 min at room temperature. The crosslinking was quenched by addition of glycine (125 mM final concentration), followed by rocking for 5 min at room temperature. Subsequently, cells were pelleted at 1000 g for 5 min, washed in PBS and pelleted again at the same speed. The final pellet was snap-frozen in liquid nitrogen and stored in −80 °C.

All subsequent steps to enrich for nuclei were carried out on ice or in 4 °C – cold room, following a protocol developed by Becker (*40*). Cell pellets were thawed and incubated for 10 min in 1 mL Hypotonic Lysis Buffer (20 mM HEPES-KOH pH 7.5, 20 mM KCl, 1 mM EDTA, 10 % glycerol, 1 % IGEPAL CA-630, 0.25 % Triton-X, 1 mM DTT, 0.2 mM PMSF, Protease Inhibitor Cocktail (cOmplete, EDTA-free, Roche #11873580001)) and disrupted with 50 strokes in a 7 mL Dounce homogenizer (Wheaton, # 357542). Nuclei were pelleted at 1,300 g for 5 min in a swinging bucket, benchtop centrifuge, washed with hypotonic buffer, and pelleted again at the same speed. Pellets were resuspended in 1 mL Nuclear Wash Buffer (10 mM Tris-HCl pH 8.0, 200 mM NaCl, 1 mM EDTA, 0.5 mM EGTA, 1 mM DTT, 0.2 mM PMSF, Protease Inhibitor Cocktail) and incubated for 10 min, rocking at 4 °C. Nuclei were pelleted at 1,300 g for 5 min and resuspended in 1.2 mL Sonication Lysis Buffer (10 mM Tris-HCl pH 8.0, 100 mM NaCl, 1 mM EDTA, 0.5 mM EGTA, 0.5 % N-lauroylsarcosine, 0.1 % sodium deoxycholate, 1 mM DTT, 0.2 mM PMSF, Protease Inhibitor Cocktail) and aliquoted into 400-μL samples (180-240 μg of DNA per sample). Sonication was carried out in 1.5 mL microtubes (Axygen®, MCT-150-C-S) using a Diagenode Bioruptor Plus UCD-300 (power HI, 20 cycles of 30 s on / 30 s off), samples were immersed in 4 °C water bath. After sonication, lysates were centrifuged at 15,000 g for 15 min at 4 °C to pellet debris. Supernatants were combined and subsequently loaded on sucrose gradients in aliquots of 300-400 μl per gradient.

#### Sucrose gradient sedimentation of chromatin

5 %–40 % linear sucrose gradients were poured into 12-mL centrifuge tubes (Beckman Coulter, #344059, 14 × 89 mm), using a Biocomp Gradient Master™. Gradients were prepared by layering bottom-up 6 mL of 5 % and 6 mL 40 % sucrose solution directly in the centrifuge tubes, capped, placed on a rotary plate on the Gradient Master and gently mixed by rotation at a 81.5° angle and a speed of 21 rpm. Sucrose solutions were prepared in 10 mM Tris-HCl, pH 8.0, supplemented with 100 mM NaCl, 1 mM EDTA, 0.5 mM EGTA, 1 % Triton-X, 0.1 % N-lauroylsarcosine, 1 mM DTT, 0.2 mM PMSF, Protease Inhibitor Cocktail. Sucrose concentrations were determined using a benchtop refractometer (Milton Roy Abbe).

Crosslinked and sheared chromatin was mixed with the 40 % sucrose solution for a 5 % final sucrose concentration and gently layered (300-400 μL, 200-360 μg of DNA) on the 5% - 40% gradient. Gradients were centrifuged using a Beckman SW 41 Ti rotor at 36,000 rpm for 4 h (4 °C), with slow acceleration and deceleration settings. Subsequently, gradients were fractionated top-down using a micropipet, generating 24 fractions of 500 μL. To remove sucrose, fractions were buffer exchanged 3-4 times into Nuclear Wash Buffer and concentrated (100 μL final volume) using 500-μL centrifugal filters with 30K cut-off (Amicon Ultra, UFC 503096). No protein or DNA loss was detected in the flow through by Bradford assay or UV-absorbance. Fractions were de-crosslinked by supplementing with SDS (1 % final concentration), transferring to PCR tubes and heating to 70 °C for at least 16 h using Peltier Thermal Cycler PTC-200 (MJ Research).

#### Western blotting

Protein concentration in fractions was quantified by Bradford assay (Bio-Rad Laboratories, #5000006). Protein samples were mixed with 4x SDS sample loading buffer (200 mM Tris-HCl, 400 mM DTT, 8 % (w/v) SDS, 30 % (w/v) glycerol, bromophenol blue) and heated to 90 °C for 10 min. Samples were loaded (equal concentration of 20-40 μg of total protein load per lane) in SDS-PAGE 4 %-20 % Tris-glycine continuous protein gels and run in SDS-containing Tris-glycine-chloride buffer (50 mM Tris, 380 mM glycine, 1 % SDS). Samples were transferred in CAPS transfer buffer (10 mM 3-[cyclohexylamino]-1-propanesulfonic acid, pH 11.3, 0.1% SDS, and 20% methanol) onto polyvinylidene fluoride membrane (PVDF) (Bio-Rad Laboratories, #1620177) for 4 h at 400 mA, 4 °C. After the transfer, PVDF membranes were washed in TBS-T buffer (20 mM Tris, pH 7.5, 150 mM NaCl, 0.1% Tween-20) and blocked in 5% nonfat diary milk in TBS-T buffer for 30 min at room temperature. Subsequently, membranes were incubated with primary antibodies overnight at 4 °C. Primary antibodies were diluted in 5% milk/TBS-T at the following concentrations: rabbit anti-CENP-A (1 μg/mL, custom), rabbit anti-CENP-C (2 μg/mL, custom), rabbit anti-H3K9me3 (2 μg/mL, Abcam, ab 8898), rabbit anti-H4-antibodies (2 μg/mL, Abcam, ab 7311) and mouse anti-Flag M2 antibodies (F1804; Sigma-Aldrich). HRP-conjugated secondary antibodies (Bio-Rad Laboratories, anti-rabbit #170-6515, anti-mouse #170-6516) were diluted 1:4,000 in 5% milk/TBS-T. Blots were developed using Super-Signal West Pico Plus Chemiluminescent Substrate (Thermo Fisher Scientific #34577) and Pierce ECL Western Blotting Substrate (Thermo Fisher Scientific #32106), visualized using a film processor SRX-101A (Konica Minolta Medical and Graphic), and analyzed with an image processor GelAnalyzer 19.1 (GelAnalyzer.com)

#### Immunofluorescence

Cells were seeded on poly-L-lysine coated glass coverslips and allowed to attach for 24 hours. Cells were then treated with or without 1 mM IAA and with or without 200 ng/mL doxycycline for 24 hours. Coverslips were then washed with PBS + 1 mM MgCl2 and CaCl2, permeabilized with PBKCl (139.7 mM KCl, 11.8 mM KH2PO4) with 0.5% Triton X-100 for 5 minutes and fixed in PBKCl/0.5% Triton X-100/3.7% formaldehyde for 10 minutes, and blocked for at least 30 minutes in antibody dilution buffer (20 mM Tris HCl, pH 7.4, 150 mM NaCl, 0.1% Triton X-100, 2% bovine serum albumin, and 0.1% sodium azide). Coverslips were then incubated for 1 hour with antibody dilution buffer containing the following primary antibodies – Transgenic CENP-N was detected using RFP antibody pre-adsorbed (α-mCherry) (Rockland Inc. Cat. No. 600-401-379) at 1 ug/ml. Centromeres were detected using α-centromere antibody derived from human CREST patient serum (Antibodies, Inc. Cat. No. 15-234-0001) at 1:100 dilution. Primary antibodies were detected using Alexa-568 or −647 conjugated goat secondary antibodies (Molecular Probes). Endogenous CENP-N was imaged directly using sfGFP fluorescence. Nuclei (DNA) were stained using 10 ug/ml Hoechst 33258.

Imaging was performed using a DeltaVision Core deconvolution microscope (Applied Precision) with a Sedat quad-pass filter set (Semrock) and monochromatic solid-state illuminators. Images were acquired using a CoolSnap HQ CCD camera (Photometrics) as Z-stacks with Z-sections of 0.2 um steps using a 60x, 1.4 NA Plan Apochromat oil immersion objective (Olympus). Image analyses were done using centromere finder (*64*), available at http://cjfuller.github.io/imageanalysistools/). Quantification of microscopy experiments involved three independent experiments with at least 100 cells per coverslip/condition per experiment. Representative images in figures were generated from maximum intensity projections of deconvolved Z-stacks using softWoRx 4.1.0 software (Applied Precision).

#### Clonogenic Survival Assay

Clonogenic survival assay was performed as described (*15*). Briefly, cells were seeded with 1000 cells/well in 6-well plates and allowed to attach for 24 hours. Cells were treated with 0.1 ug/ml doxycycline and/or 0.1 mM IAA before fixing and staining with Crystal Violet solution (0.25% crystal violet in 3.5% formaldehyde and 72% methanol) for 30 minutes. Cells were then destained in water and allowed to air dry before being imaged (using a scanner).

#### In Silico System Construction

Using the 6EQT initial coordinate structure (*13*) and the nucleosome dimer cryo-EM map, we created models for use in molecular dynamics simulations to understand the influence of CENP-N on the nucleosome dimer. We modeled the missing loops and tails in each system by using Modeller via the Chimera graphical user interface (*65*) (*66*). This resulted in the histone tails extended outward from each nucleosome with unique intrinsically disordered conformations. Once structures were assembled, the coordinates were aligned to the cryo-EM map using the Molecular Dynamics Flexible Fitting (MDFF) method within the NAMD 2.13 engine in implicit solvent (*67*). After 300 ps of biasing, the system reached an equilibrium and could not be refined further. The structure from this point was used to seed simulations of the dimer containing one of the following: 2 CENP-N, 1 CENP-N, or no CENP-N. The latter two systems were created by removing the CENP-N coordinates from the initial structure file.

#### Simulation Methods

All systems were prepared with tleap from the AmberTools18 software package (*68*)). Each system was solvated in a TIP3P water box extending at least 10 Å from the solute (*69, 70*). Using Joung-Cheatham ions, the solvent contained 150 mM NaCl, and sodium cations to neutralize negative charges (*71*). The AMBER14SB and BSC1 force fields were used for protein and DNA interactions, respectively (*72*). All simulations were performed locally using the Amber18 engine on GeForce GTX 1080 GPUs via the Pinot computing cluster (*68*).

Systems were minimized twice for 10,000 steps, first with a 10 kcal·mol-1·Å-2 harmonic restraint applied to the solute and then followed by no restraints. Using the Langevin thermostat and an electrostatic cutoff distance of 10 Å with long range interactions treated with particle mesh Ewald calculations (*73*), systems were then heated in the NVT ensemble from a temperature of 5 K to 300 K over 5 ps with a 10 kcal/mol kcal·mol-1·Å-2 solute restraint (*73*). This restraint was then reduced from 10 kcal/mol to 0 kcal/mol over 600 picoseconds in the NPT ensemble, followed by a 1 ns equilibration period to assess the system stability. Production runs with no restraints and a temperature of 300 K were also performed in the NPT ensemble using the Berendsen barostat. To allow for the extended timestep of 4 femptoseconds, we used Hydrogen Mass Repartitioning (HMR) which increases the mass of hydrogens while equally decreasing the parent atom masses to moderate the vibrations of high-frequency bonds (*74–76*). Simulations required 100 ns of equilibration time to allow the tails to relax down onto the DNA of their respective nucleosomes. To boost the speed and overall efficiency, after equilibration, the solvent was removed from each system and then resolvated. Since the tails had collapsed, the overall dimensions of the solute were reduced translating to less water required upon solvation, resulting in a fewer number of particles and a boost to the overall simulation speed. Simulations of each system were performed in triplicate for 600 ns each, totaling in 5.40 μs. Allotting for the 100 ns mentioned above, we have amassed 4.50 μs of post-equilibration data.

Movies of the stacked-nucleosome simulations were created by using a combination of Visual Molecular Dynamics (VMD; (*77*) and FFmpeg (*78*). VMD was used to generate the individual frames which employed Tachyon ray tracing (*79*) to implement advanced rendering features such as ambient occlusion lighting. The frames were merged to produce the final video by using FFmpeg, a free and open-source command-line based video processing software (*78*).

#### Contact Analysis

Analysis was conducted using AmberTools18 from the AMBER18 software package (*68*). Protein-DNA contacts were defined between the heavy atoms of residues within 4.0 Å of one another. The statistical inefficiency was calculated for each residue to determine the number of statically independent data points throughout each simulation.

### Supplementary Text

#### 2.68 Å structure of CENP-A α satellite nucleosome in complex with CENP-N^NT^

The structures of CENP-N^NT^ in complex with CENP-A nucleosome containing the 601 nucleosome positioning sequence were reported by multiple labs including our own (*13, 14, 19*). The structure of a complex consisting of CENP-N^NT^, CENP-C^CC^ and CENP-A nucleosomes reconstituted on α satellite DNA was also solved (*16*). In these structures, the resolution for the CENP-A nucleosome ranged from 3.5-4.0 Å, while CENP-N^NT^ was at 4.0-4.5 Å resolution. The side chains of CENP-N were not resolved in the density. In our new map, the resolution of CENP-N was higher than 3.5 Å (Fig S1C), revealing density for key CENP-N amino acid side chains that are involved in the interaction with CENP-A (fig. S14B), and largely confirming the accuracy of our previous structure (6C0W).

The resolution of the nucleosome (reconstituted with a 147 bp palindromic DNA fragment derived from α-satellite DNA, as described in (*62*), pdb 1kx5) in our new map is ∼2.8-3.0 Å (fig S1C, D), and reveals density for individual base-pairs of nucleosomal DNA. This allowed us to precisely position each nucleotide for the majority of the DNA (Fig. S14D). When attempting to fit the DNA derived from 1kx5 into the map, we observed a staggered 1 bp compression of DNA around SHL ±2 (bp 12-20; central ‘dyad’ base pair designated as 0, fig S14E). This suggests that the ‘natural’ length of DNA organized by α-satellite DNA (and likely also by other DNA) is 145 bp rather than 147 bp as suggested by the crystal structures. The compression of DNA observed in the crystal structure is caused by the end-to-end stacking of DNA in the crystal lattice (*80*), and further suggests that the ‘stretching’ of DNA required for crystal lattice formation with 145 bp DNA (*81*) is the ‘relaxed’, natural in-solution state for most nucleosomal DNA sequences. Of note, the resolution of our earlier structure did not allow us to address this issue with 601 DNA in 6C0w. Our maps further confirm SHL ±2 as a ‘soft-spot’ to absorb DNA length variability (*82*).

**Fig. S1.**
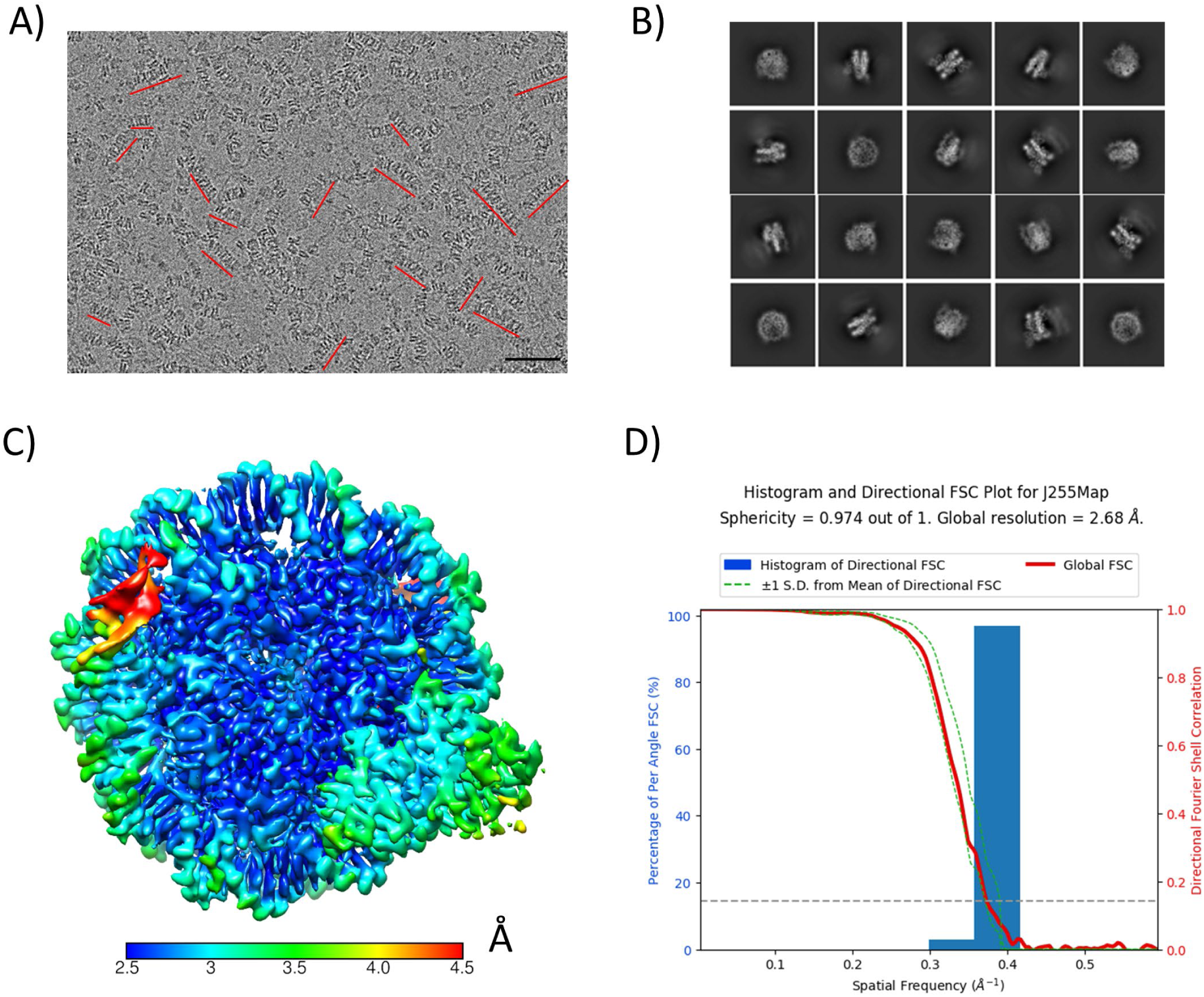
CryoEM analysis of CENP-N in complex with CENP-A mono-nucleosomes (reconstituted with 147 bp palindromic α satellite DNA). **A)** Raw cryoEM micrograph. Stacks are indicated by red lines. Size bar is 50 nm. **B)** 2D classifications of particles at the mono-nucleosome level. **C)** Local resolution of the 3D map for CENP-N in complex with CENP-A mono-nucleosomes. The color key indicates the resolution. **D)** 3DFSC analysis of reconstructed 3D map.

**Fig. S2.**
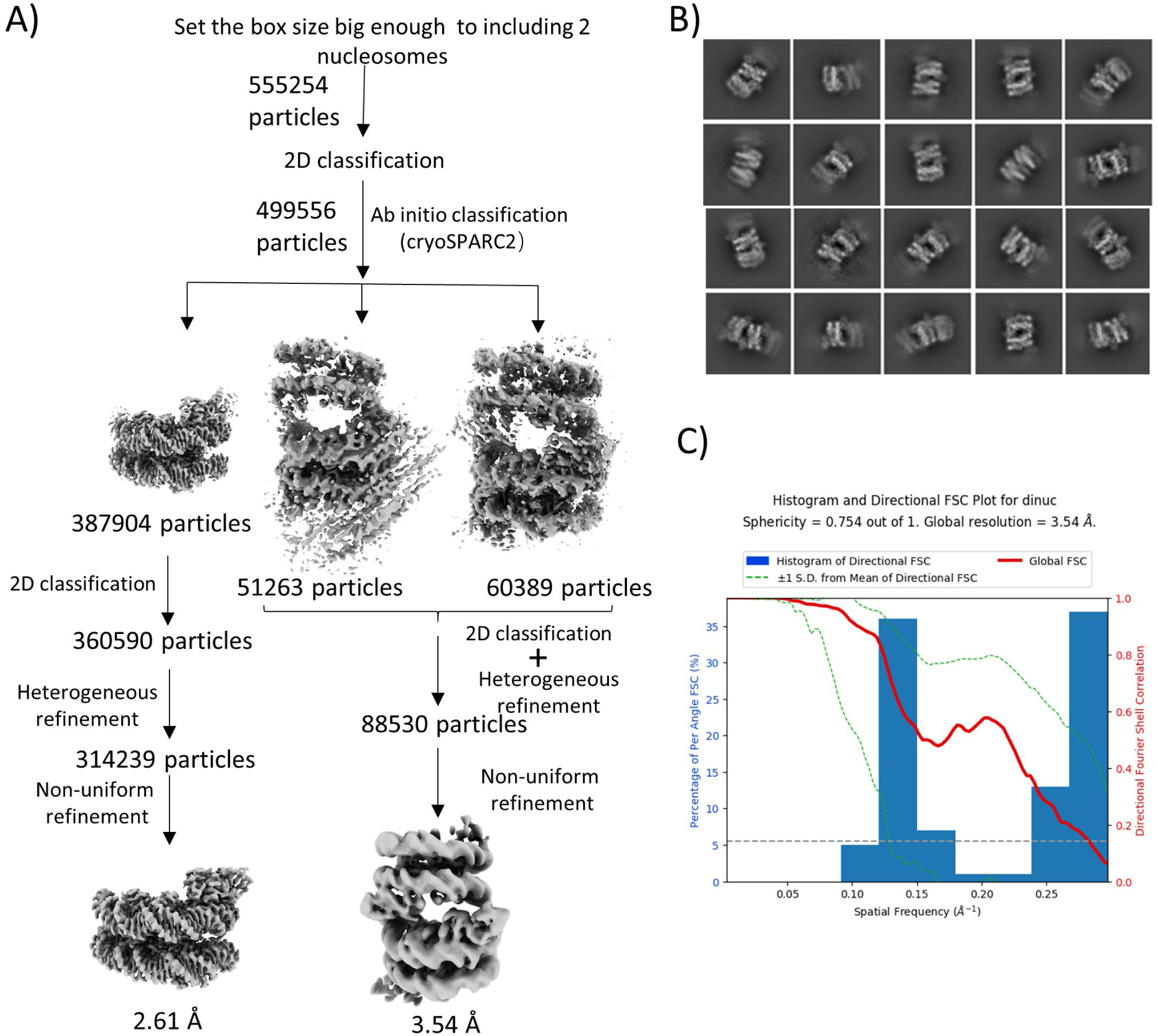
CryoEM analysis of CENP-N mediated stacks of CENP-A mono-nucleosomes (reconstituted with palindromic α-satellite DNA. **A)** Flow-chart on the analysis of di-nucleosomes in the stacks. **B)** 2D classification of the di-nucleosome with CENP-N. **C)** 3DFSC analysis of the reconstructed 3D map.

**Fig. S3.**
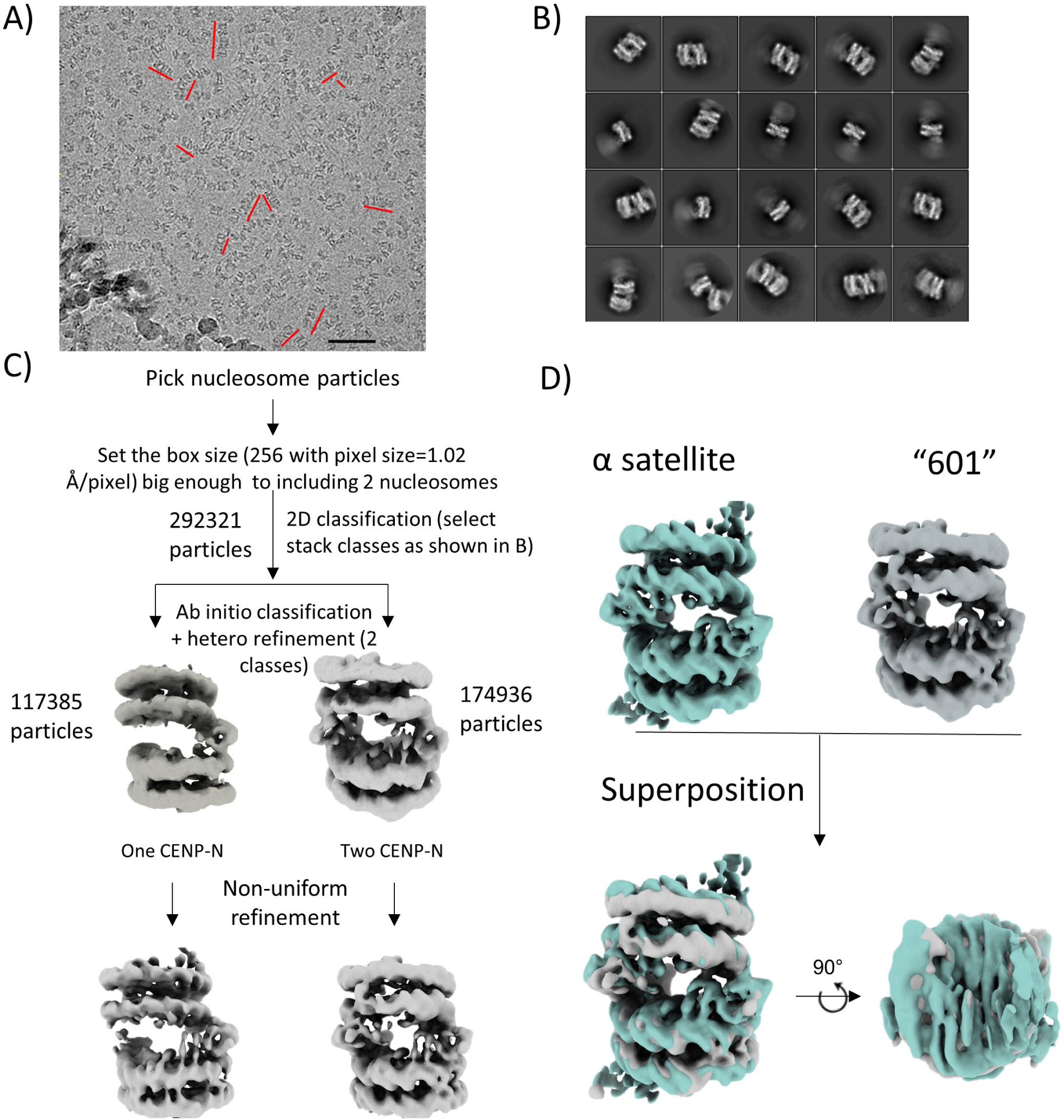
CryoEM analysis of CENP-N mediated stacks of CENP-A mono-nucleosomes reconstituted with 601 DNA (*21*). **A)** Representative cryoEM micrograph. Stacks are highlighted by red lines. Size bar is 50 nm**. B)** 2D classification of the di-nucleosome with CENP-N in the stacks. **C)** Flow-chart on the analysis of di-nucleosomes in the stacks. **D)** Comparison of di-nucleosome maps derived for α satellite and 601 CENP-A nucleosomes.

**Fig. S4.**
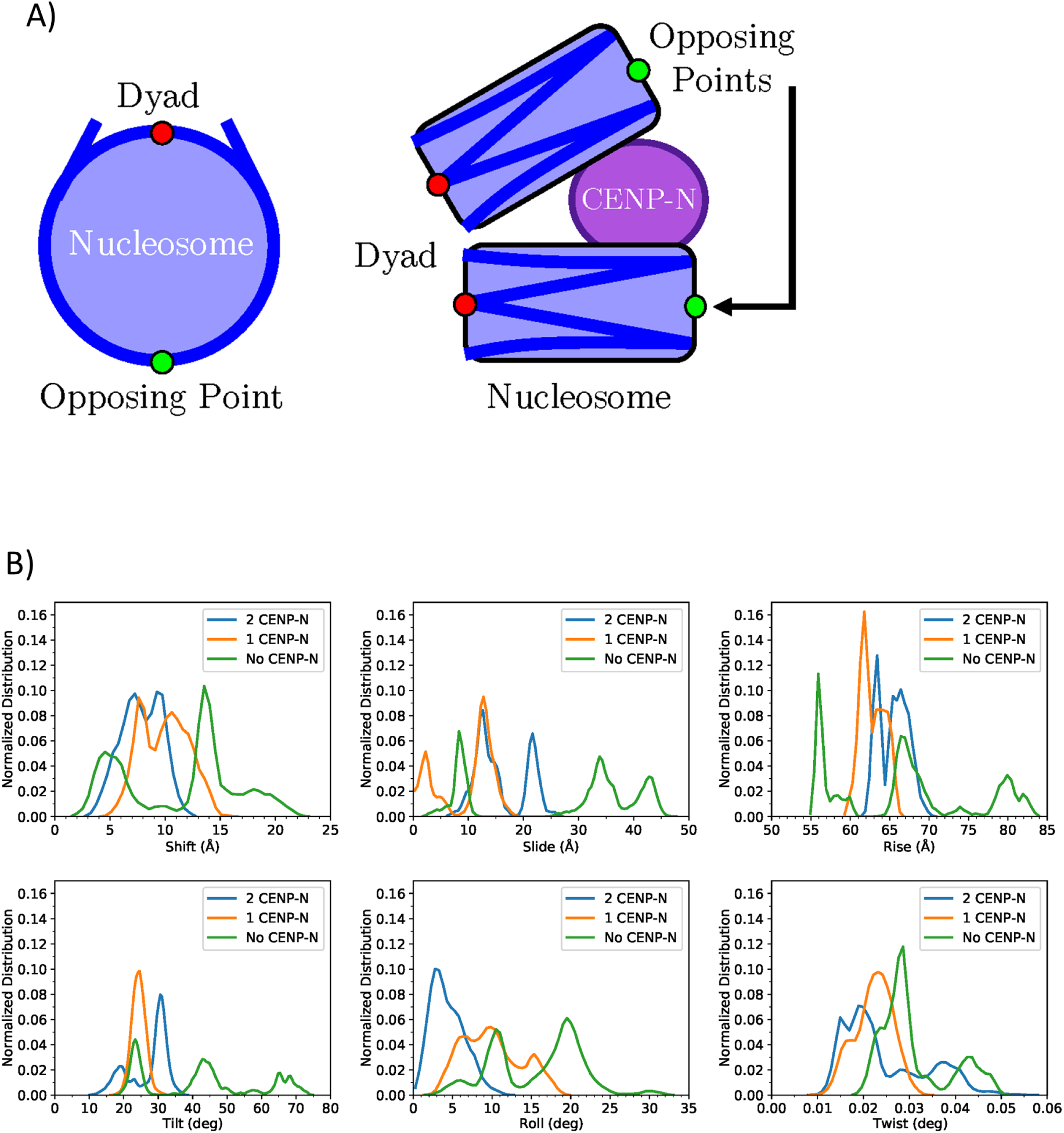
MD simulations of stacked nucleosomes with 0, 1, or 2 CENP-N. **A)** Diagram of the points used to construct stacked-nucleosome sampling graphs as depicted in Figure 1C, in addition to nucleosome parameters calculated for stacked-nucleosome simulations. Nucleosomes (blue) are shown in face-on (left) and profile (right) viewpoints, with DNA represented in a darker blue. Dyad points and their opposing points are represented as small red and green circles, respectively. CENP-N is shown in purple to provide a point-of-reference. **B)** From left to right, six histograms of stacking parameters for di-nucleosome systems: Shift, Slide, Rise (top), and Tilt, Roll, Twist (bottom), in analogy to the parameters used to describe the geometry of the DNA double helix. Histograms do not contain the first 100 ns of simulation time which was allotted for each system to achieve equilibration.

**Fig. S5.**
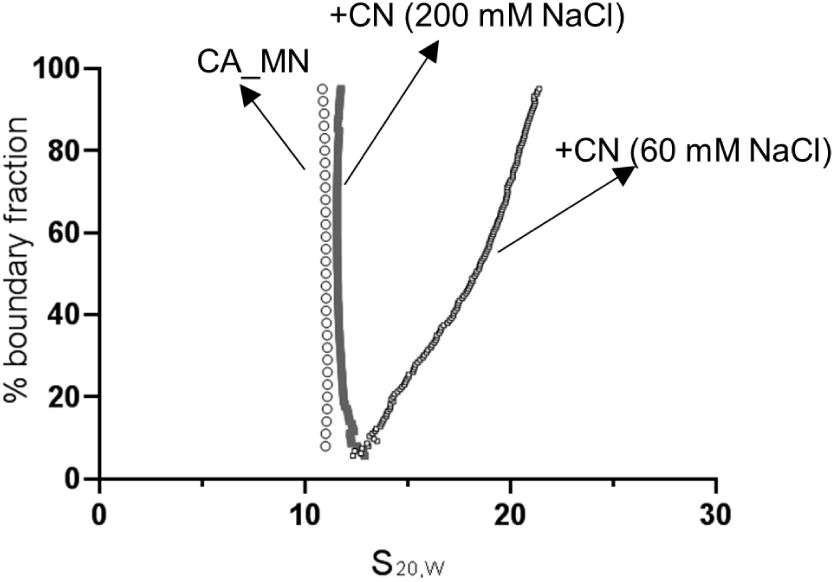
Salt concentration affects nucleosome stack formation. AUC analysis (van Holde-Weischet plots) of CENP-N in complex with CENP-A mono-nucleosomes at 60 and 200 mM NaCl). CN: CENP-N^1-289^ in complex with CENP-A mono-nucleosome. CA_MN: CENP-A mono-nucleosome.

**Fig. S6.**
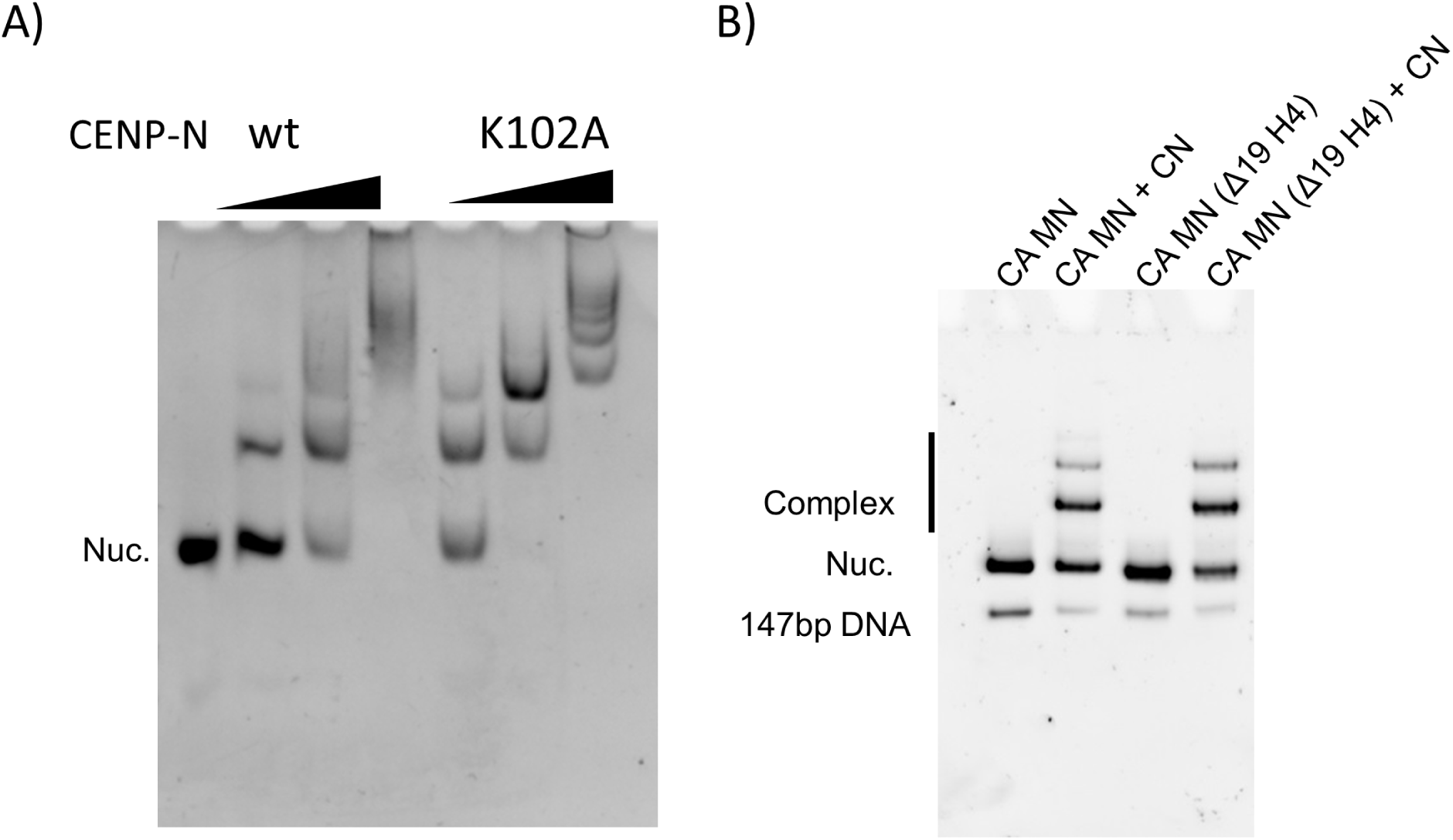
Contributions of CENP-N and H4 N-terminal tail to complex formation. **A)** CENP-N mutant (K102A) binds to CENP-A nucleosomes as well as wild-type CENP-N (5% native PAGE). 250 nM CENP-A nucleosome was combined with CENP-N at ratios of 2:1, 4:1 and 8:1 in buffer containing 50 mM NaCl, 20 mM Tris-HCl (pH 7.8), 1 mM EDTA, 1 mM DTT. **B)** deletion of the H4 N-terminal tail (Δ19) does not affect the specific interaction between CENP-N and CENP-A nucleosomes. CENP-N was mixed with CENP-A nucleosome containing full length H4 or (Δ19) H4 at a 2:1 ratio in the same buffer as in A).

**Fig. S7.**
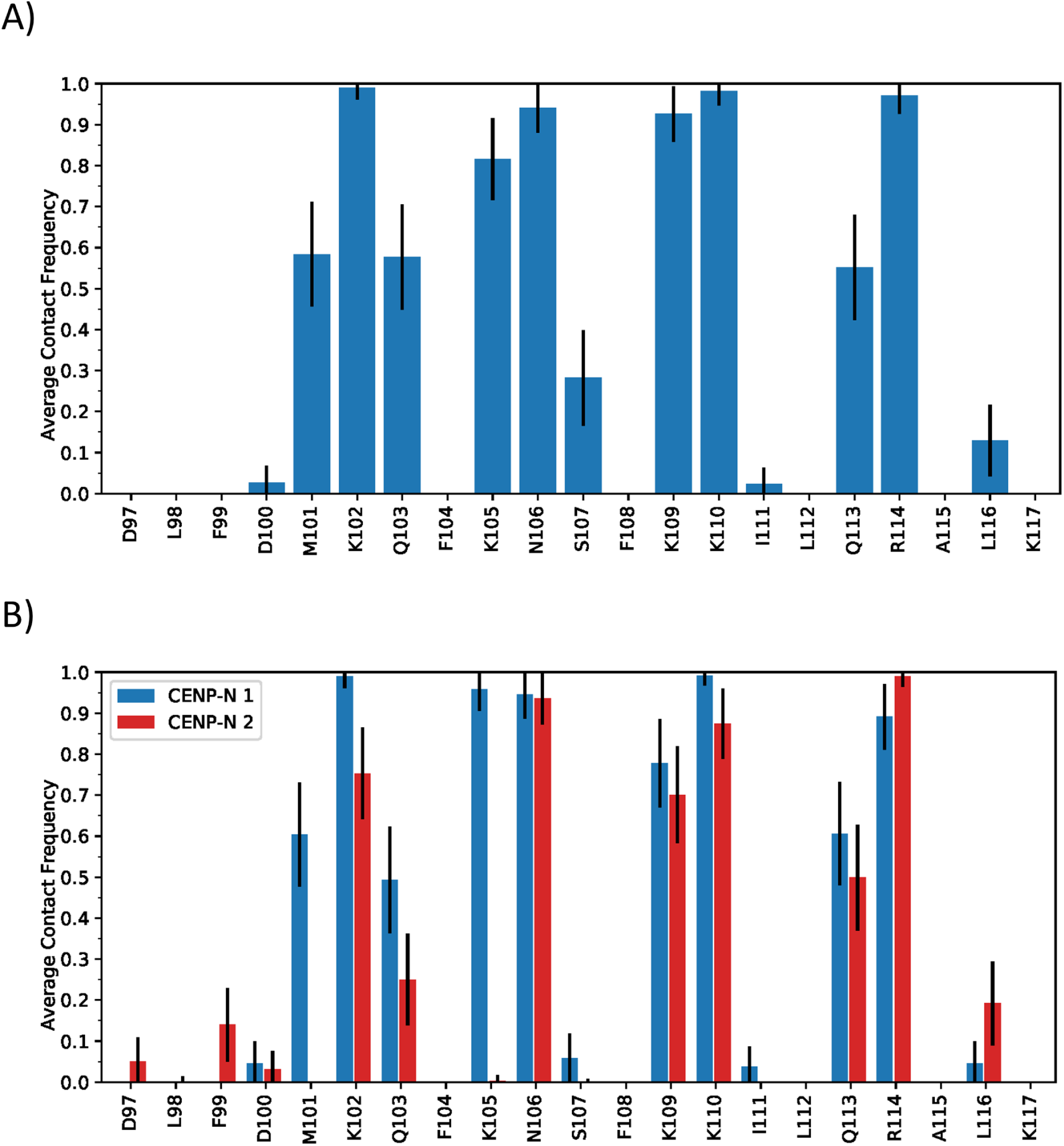
Residue contacts of the CENP-N α6-helix with DNA of the DNA-directed nucleosome from simulations containing one (A) and two CENP-N (B). CENP-N 1 and CENP-N 2 are distinguished by the binding orientation of the α6-helix with the DNA grooves. CENP-N 1 (blue) binds directly into the DNA minor groove while CENP-2 (red) does not. Protein-DNA contacts were defined between the heavy atoms of residues within 4.0 Å of one another.

**Fig. S8.**
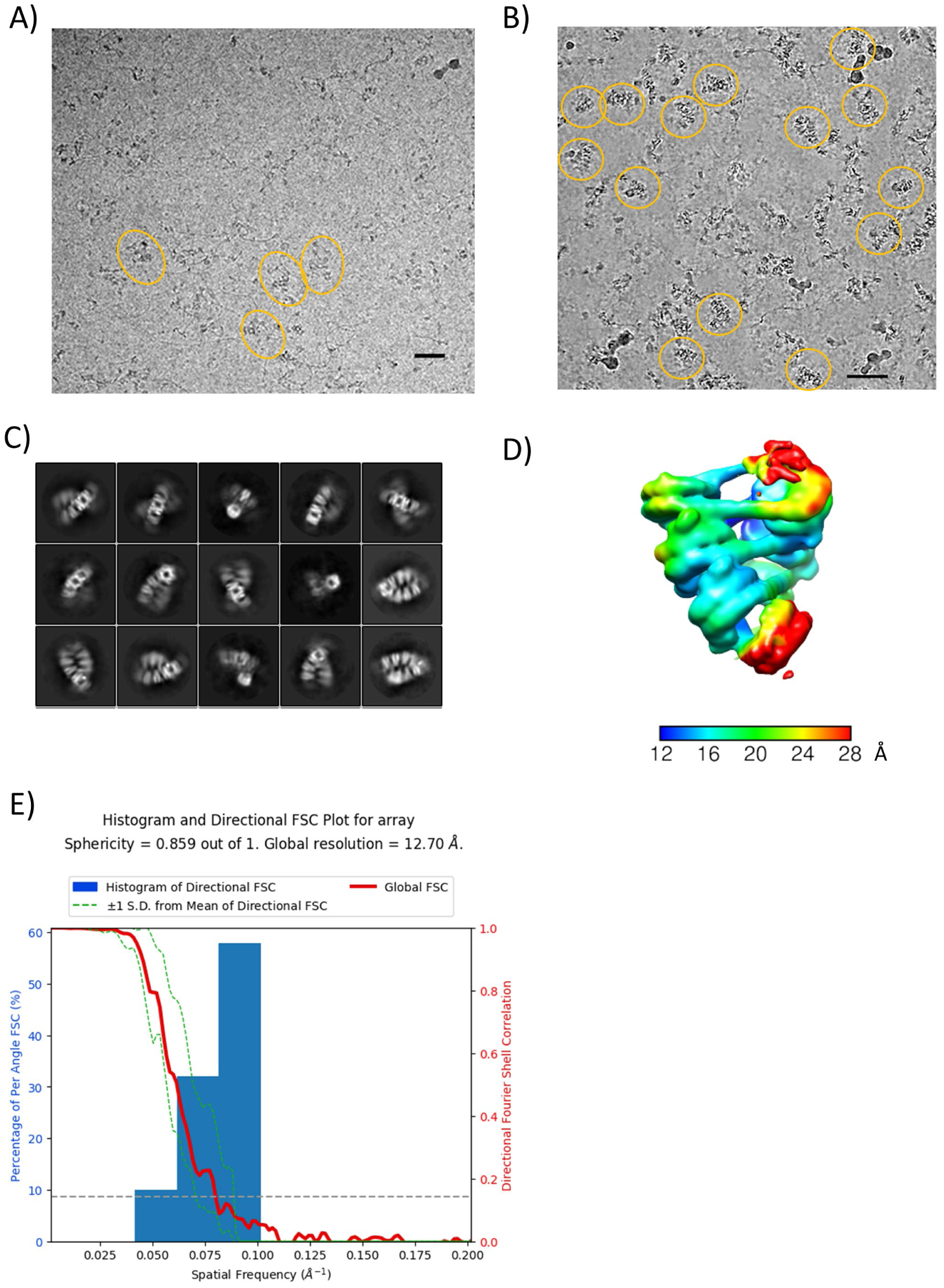
CryoEM analysis of 12mer nucleosomal arrays in presence of CENP-N. **A)** raw cryoEM image of CENP-N in complex with 12-207mer 601 array. Size bar is 50 nm. **B)** raw cryoEM image of CENP-N in complex with 12-167mer array. Size bar is 50 nm. **C)** 2D classification of CENP-N in complex with 12-167mer array. **D)** Local resolution map of chromatin fiber. Color key indicates the resolution. **E)** 3DFSC analysis of cryoEM electron map.

**Fig. S9.**
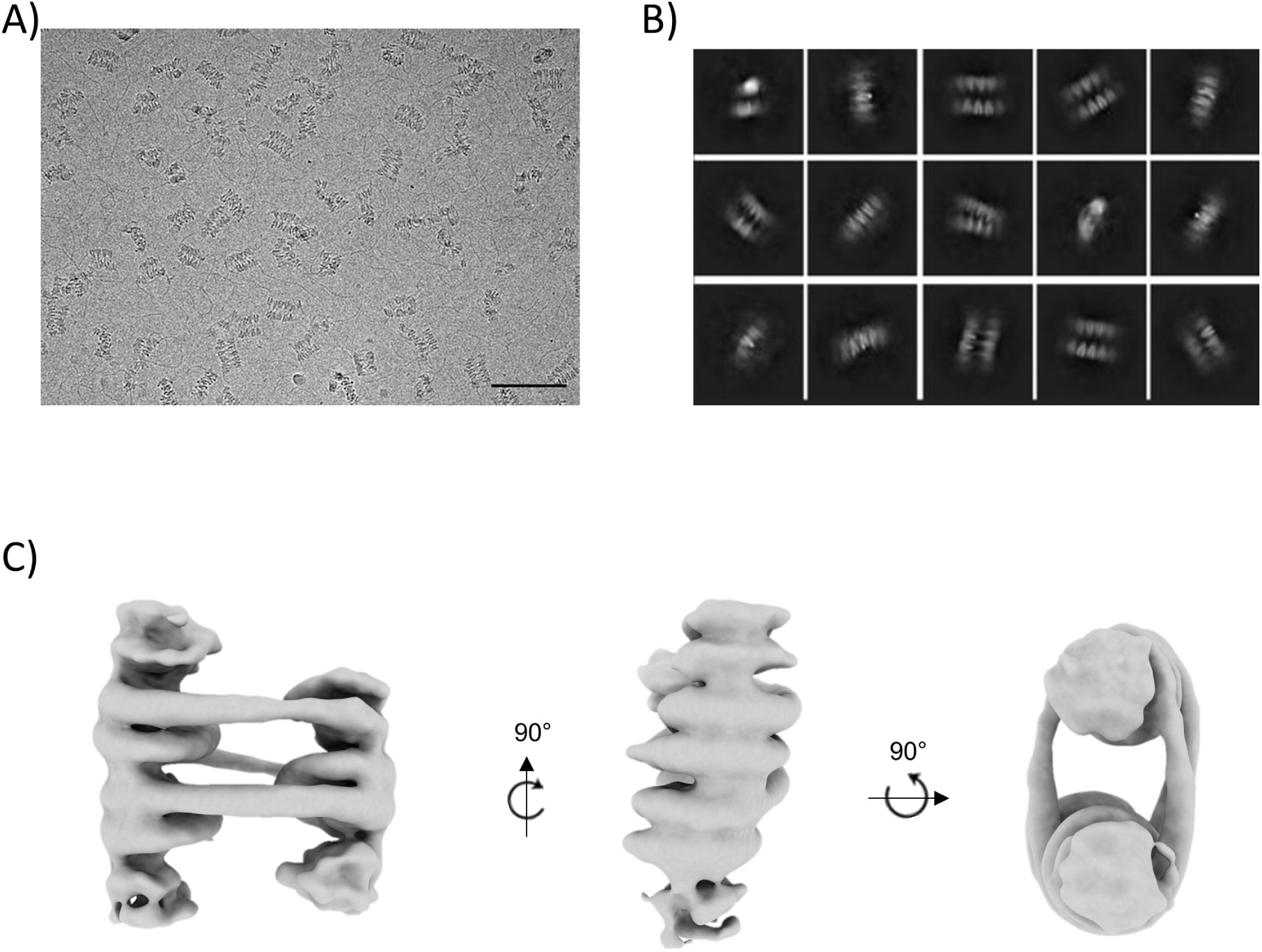
CryoEM analysis of crosslinked chromatin arrays without CENP-N. **A)** raw cryoEM image. Size bar is 100 nm **B)** 2D classification of CENP-N in complex with 12-167mer array. **C)** Low resolution 3D cryoEM map illustrates ladder-like arrangement of nucleosomes.

**Fig. S10.**
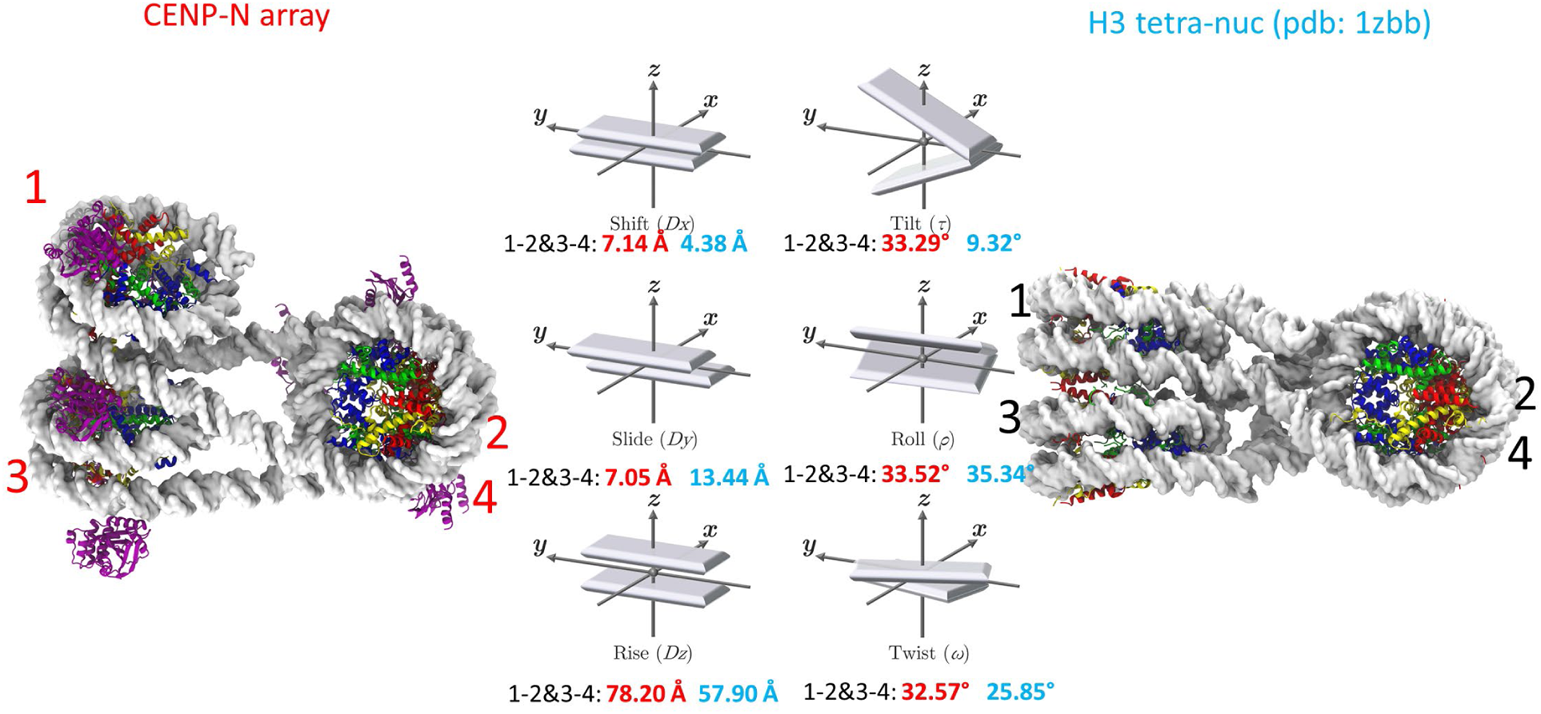
Comparison between tetra-nucleosomes in a chromatin fiber with CENP-N, and of a canonical tetra-nucleosome. Analysis of the relative orientation of the nucleosomes, in analogy to DNA base pair analysis, is shown in the middle panel, with numbers for the CENP-N array in red, and tetranucleosome array in black.

**Fig. S11.**
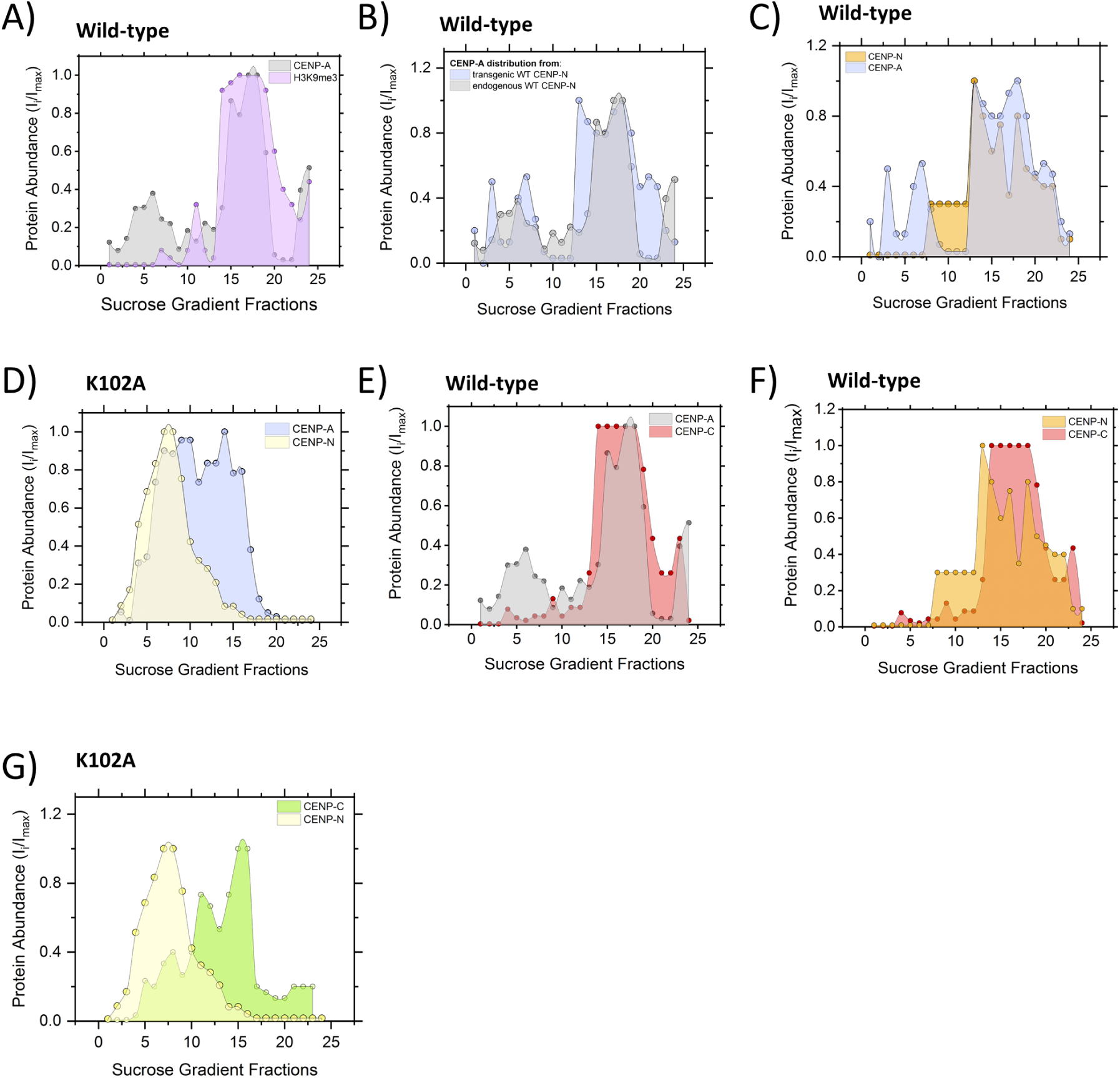
Western blot detection of centromeric proteins across 5-40% sucrose gradients. **A)** Comparision of CENP-A (gray) and H3K9me3 (lilac) distribution in the presence of endogenous WT-CENP-N. **B)** CENP-A distribution in the presence of endogenous (gray) and transgenic (blue) WT-CENP-N. **C)** Comparision of CENP-A (blue) and CENP-N (orange) in the presence of WT CENP-N. **D)** Comparision of CENP-A (blue) and CENP-N (yellow) in the presence of K102A CENP-N. **E).** Comparision of CENP-A (gray) and CENP-C (red) distribution in the presence of endogenous WT-CENP-N. **F)** CENP-N (orange) and CENP-C (red) distribution in the presence of endogenous WT-CENP-N **G**) CENP-N (yellow) and CENP-C (green) distribution in the presence of K102A CENP-N.

**Fig. S12:**
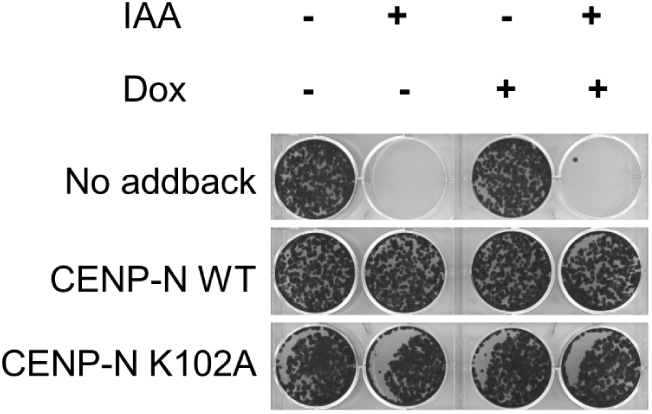
Clonogenic survival assay showing CENP-N AID cells with the indicated transgenic CENP-N variant treated with or without 0.1 mM IAA and doxycycline for 14 days. After treatment of 1000 seeded cells/well, surviving colonies were fixed and stained with crystal violet stain.

**Fig. S13.**
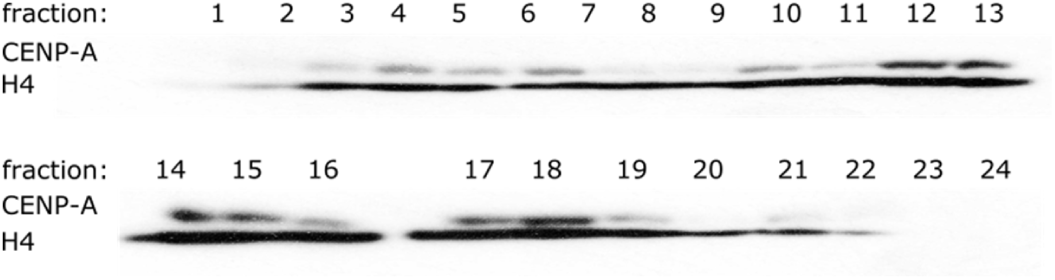
Western blot of CENP-A and H4 histone distribution across 5-40 % sucrose gradient. Presence of CENP-A signal indicates fractions containing the centromeric chromatin whereas H4 signal represents the overall amount of chromatin loaded onto a SDS gel.

**Fig. S14.**
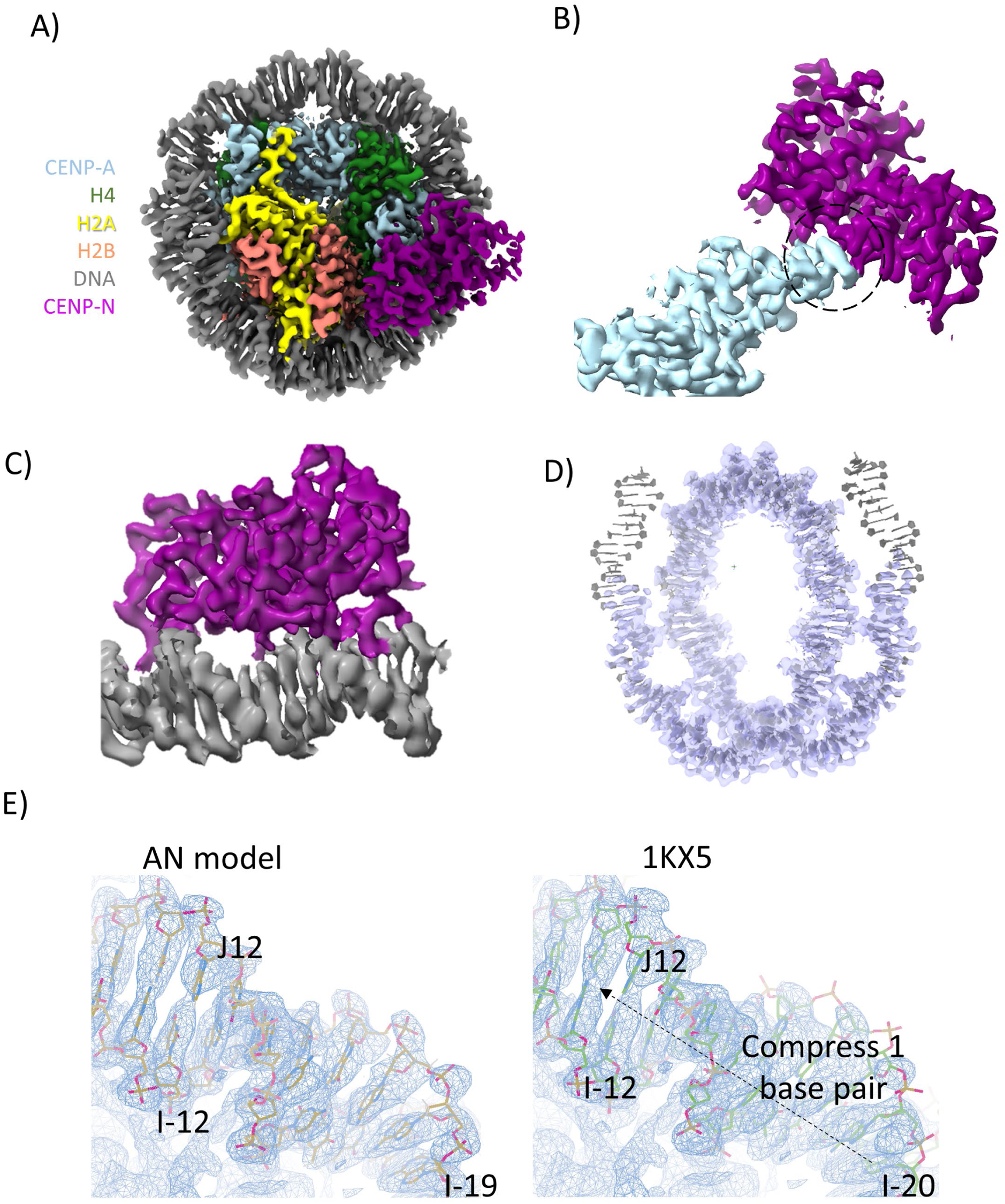
The structure of CENP-A α satellite nucleosome in complex with CENP-N^NT^. **A)** 2.65 Å cryoEM map of CENP-A α satellite nucleosome in complex with CENP-N^NT^. The components are colored as indicated. **B)** The density of the interface between CENP-N and histones. **C)** The density of the interface between CENP-N and nearby DNA. **D)** The model to density of DNA. **E)** α satellite DNA is compressed by one base pair in the crystal structure of nucleosome, compared to the cryoEM structure where DNA ends are unconstrained.

**Movie S1**. Simulation of the stacked-nucleosomes bound to two CENP-N. Shown is the DNA (white), CENP-N (purple), histone H3 (blue), histone H4 (green), histone H2A (yellow), and histone H2B (red). The simulation time evolution is presented in the bottom right-hand corner of the video.

**Movie S2**. Simulation of the stacked-nucleosomes bound to one CENP-N. Shown is the DNA (white), CENP-N (purple), histone H3 (blue), histone H4 (green), histone H2A (yellow), and histone H2B (red). The simulation time evolution is presented in the bottom right-hand corner of the video.

**Movie S3.** Simulation of the stacked-nucleosomes not bound to any CENP-N. Shown is the DNA (white), histone H3 (blue), histone H4 (green), histone H2A (yellow), and histone H2B (red). The simulation time evolution is presented in the bottom right-hand corner of the video.

## Notes

### Competing Interest Statement

The authors have declared no competing interest.

